# Cell type-specific proximity labeling of organ secretomes reveals energy balance-dependent proteomic remodeling

**DOI:** 10.64898/2026.01.11.698831

**Authors:** Kaja Plucińska, Charlotte R. Wayne, Henry Sanford, Boby Mathew, Nathalie Ropek, Stephanie M. Adaniya, Corey Model, Nicolás Gómez-Banoy, Ksenia Morozova, Xiongwen Cao, Jeffrey M. Friedman, Ken H. Loh, Paul Cohen, Ekaterina V. Vinogradova

**Affiliations:** Laboratory of Molecular Metabolism, The Rockefeller University, 1230 York Avenue, New York, NY 10065, USA; Laboratory of Chemical Immunology and Proteomics, The Rockefeller University, 1230 York Avenue, New York, NY 10065, USA; Department of Comparative Medicine, Yale University, School of Medicine, New Haven, CT 06520, USA; Department of Biochemistry, University of Washington, Seattle, WA 98195, USA; Department of Chemistry, Stanford University, Stanford, CA 94305, USA; Division of Endocrinology, Department of Medicine, Memorial Sloan Kettering Cancer Center, New York, New York 10065, USA; Division of Endocrinology, Diabetes and Metabolism, Department of Medicine, Weill Cornell Medicine, New York, New York 10065, USA; Current position: Shanghai Key Laboratory of Regulatory Biology, Institute of Biomedical Sciences, School of Life Sciences, East China Normal University, Shanghai 200241, China; Laboratory of Molecular Genetics, The Rockefeller University, 1230 York Avenue, New York, NY 10065, USA

**Keywords:** Proximity labeling, TurboID, obesity, inflammation, ER proteomics, plasma proteomics

## Abstract

Intercellular communication is critical for maintaining organismal metabolic homeostasis. Here, we present a new method enabling temporally controlled, cell type-specific labeling of secreted and membrane proteins in key metabolic tissues. The method employs a genetically encoded proximity-labeling strategy by targeting a Cre-dependent TurboID ligase to the endoplasmic reticulum (ER) in ES cell-derived mice. Expression of TurboID in liver, adipose tissue, and spleen enabled the characterization of organ-specific ER proteomes at baseline and in response to fasting, inflammation, and dietary obesity, revealing tissue-and perturbation-specific changes and augmenting our understanding of how the proteomes of individual tissues change to regulate systemic energy balance. This comprehensive resource represents an important advance toward understanding both how cell-to-cell communication changes in response to energy homeostasis and how it contributes to these alterations. This method is broadly applicable and provides a means for identifying biomarkers and therapeutic targets across a wide range of tissues.

## INTRODUCTION

In multicellular organisms, the maintenance of energy homeostasis requires the orchestration of metabolic processes across and between numerous cell types and tissues. This complex coordination is mediated by secreted factors that can act locally or at a distance to regulate processes including switching between anabolic and catabolic states and linking nutritional status with other vital functions such as immunity. Autocrine and paracrine factors modulate tissue development and remodeling, while endocrine mediators such as insulin, glucagon,^1^ and leptin^2^ are pivotal for regulating systemic glucose homeostasis, food intake, and energy balance.^3–5^ Moreover temporal differences in their expression (i.e. acute vs. chronic changes) have distinct effects within and outside cells including alterations of metabolic flux, signal transduction, endoplasmic reticulum (ER) stress, and energy balance among many others.

Despite extensive studies, the full extent of the molecular mechanisms that coordinate systemic responses to positive and negative energy balance remains incomplete. During negative energy balance, induced by fasting or systemic inflammation, endocrine and paracrine networks reprogram metabolism to sustain ATP production and preserve glucose for essential tissues but we still lack an understanding of the complete basis by which organ-specific secretomes regulate metabolism and immune function.^6,7^ Conversely, positive energy balance driven by chronic nutrient excess activates insulin and mTOR signaling to promote anabolic metabolism and lipid storage, but also drives ER stress, mitochondrial dysfunction, and inflammation. How these pathways interact across tissues from adaptive to maladaptive states also remains unresolved. We hypothesized that systematic mapping of tissue- and cell type-specific secretomes in multiple key tissues across energy states would reveal known and novel molecular mediators that respond to and regulate metabolic and immune signaling.

The creation of such an organ-specific atlas of secreted proteins across metabolic states has been a longstanding goal with the potential to provide insights into both normal physiology and disease mechanisms. However, the creation of an atlas of secreted proteins has been limited by significant technical challenges. First, circulating proteins exhibit a dynamic range of ten orders of magnitude in concentration, making it particularly difficult to identify low abundance proteins (i.e. in the ng/mL range or lower), which is the level at which most hormones circulate.^8^ Additionally, tracing the source of a secreted protein in blood or other body fluids is not possible with traditional mass spectrometry because it does not provide information on cell type specificity, nor does it provide insight into paracrine versus endocrine factors. While inferences can be gained from transcriptomic profiling, RNA levels do not always predict secreted protein levels, as they do not account for post-transcriptional and post-translational regulation or secretory dynamics.^9^ Furthermore the use of *in vitro* models to identify secreted proteins is limited as they do not fully recapitulate *in vivo* biology and are limited to cell types that can be cultured with fidelity.

The use of biorthogonal amino acid tagging (BONCAT)^10^ or proximity labeling (PL)^11^ has substantially enhanced our ability to map cell-type selective proteomes by providing precise information on the origin, identity, and spatiotemporal dynamics of intracellular and secreted proteins.^12^ PL catalyzes the proximity-dependent modification of proteins, which can be achieved by the introduction of engineered versions of biotin ligase BirA, such as R118G BirA (BioID) and BirA*G3,^13,14^ and TurboID,^11,15^ each of which promiscuously biotinylates resident or transiently associated proteins in a subcellular compartment in a cell type-specific manner. Further specificity can be achieved by targeting the enzyme to a particular cell type using conditional genetic systems. The biotinylated proteome can then be enriched with streptavidin (SA) beads and identified via mass spectrometry (MS).^11,12,15^ This was elegantly demonstrated in Drosophila using a muscle-specific ER-anchored BirA*G3 to label classically secreted proteins in response to exercise^14^ and in mice via AAV-mediated expression of an ER-linked BioID^16^ or TurboID fused with Ces61b (ER membrane)^17^ or KDEL (ER lumen). The use of these AAVs has enabled the characterization of secreted proteomes from myeloid cells under various physiologic states *in vivo*.^18,19^ However, the use of AAV can be problematic because of the typical artifacts from incomplete or off-target viral transduction and the inflammatory response provoked by viral infection. These limitations can be overcome by an introduction of Cre-dependent expression of TurboID targeted to the secretory pathway *in vivo*. We thus set out to generate a mouse line that could be used to profile secreted proteins in a cell type-specific manner.

Here we report the development and validation of a genetically encoded proximity labeling TurboID^KDEL^ mouse for cell type-selective labeling of ER-resident or secreted proteomes *in vivo*. We used this approach to create an atlas of ER-resident and secreted proteomic signatures in two different depots of white adipocytes, hepatocytes, and B cells at baseline and after fasting, inflammation, and early and advanced diet-induced obesity. Validation of the method allowed us to generate comprehensive, high-resolution data characterization of the levels of ER-localized and secreted proteins, providing new and potentially critical insights into the molecular networks governing energy balance. This included the identification of novel cell-type enriched circulating mediators linked to changes in energy balance with potential relevance to human health and disease.

## RESULTS

### Cell type-specific ER-proximity labeling

To enable *in vivo* cell type-specific proximity labeling of the secretory pathway proteome in a non-immunogenic system, we developed a knock-in mouse line in which TurboID fused to an ER retention signal (KDEL) and a V5 epitope tag was targeted to the Rosa26 locus (Figure 1A). An upstream LoxP-flanked STOP cassette restricts expression to Cre-expressing cells, enabling cell-specific expression. We crossed this TurboID^KDEL^ line to mice expressing Cre recombinase under the control of cell type-specific promoters, including Adiponectin-Cre,^20^ Albumin-Cre,^21^ and CD19-Cre^22^ for selective expression in adipocytes, hepatocytes, and B lymphocytes (Figure 1B). Biotin supplementation in drinking water led to robust biotinylation in tissues and plasma (Figures 1B-1C, S1A-S1C), while intraperitoneal injection produced weaker and less uniform labeling (Figure S1D). The specificity of TurboID expression was confirmed by SA-HRP and V5-tag Western blotting (Figures 1C, S1A-S1C) and SA immunohistochemistry (Figure 1D). Cre-negative controls showed only a background signal of endogenous biotinylated proteins,^11^ and distribution of biotin-labeled proteins was consistent with cell-type specific TurboID expression with limited signal from other tissues not expressing Cre (Figures S1A-S1C).

**Figure 1.**
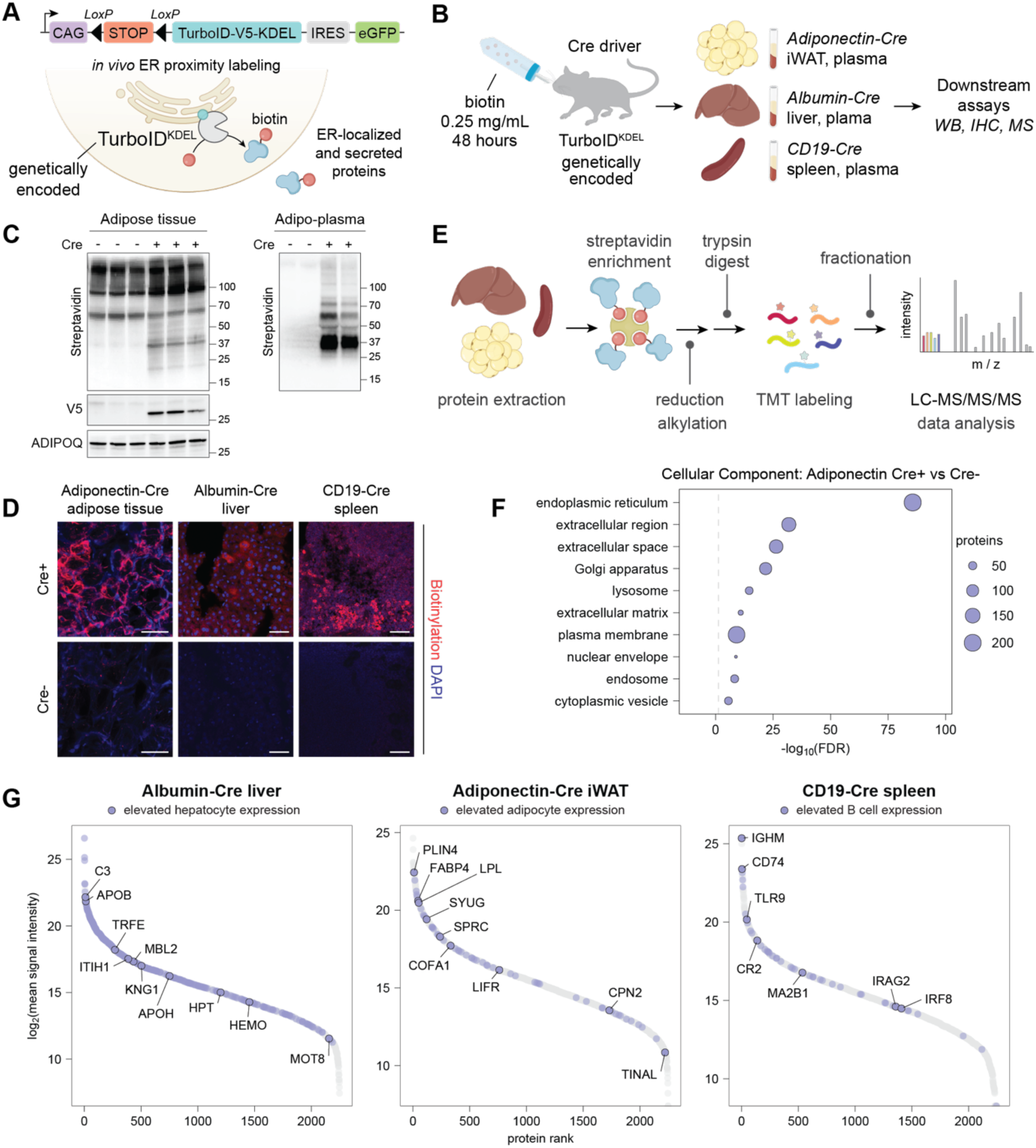
Introduction to TurboID workflow and validation of tissue-specific *in vivo* proximity labeling. (A) Schematic of LoxP-flanked TurboID-KDEL construct and ER-localized TurboID proximity labeling system. (B) Experimental design for comparison of three tissues under basal conditions. (C) Western blot showing TurboID-catalyzed biotinylation of inguinal adipose tissue and plasma from biotin-treated Adipo-TurboID^KDEL^ mice, as well as V5 tag indicative of TurboID expression. (D) Immunohistochemistry images of inguinal white adipose tissue (iWAT), liver, and spleen tissue in TurboID^KDEL^ mice with and without expression of Adiponectin-Cre, Albumin-Cre or CD19-Cre. Biotinylation is shown in red (streptavidin for iWAT, neutravidin for liver and spleen). Scale bar 50 µm. (E) Workflow for tissue processing, affinity purification, and analysis by Tandem-Mass-Tag (TMT) mass spectrometry. (F) Pathway analysis illustrating predicted subcellular localization of proteins enriched in basal Adipo-TurboID^KDEL^ Cre+ relative to Cre- samples. Dashed line indicates false discovery rate (FDR) of 5%. Gene Ontology term enrichment analysis was performed with GO Slim Cellular Component with the full mouse genome used as the background list. (G) Protein abundance plots highlighting cell-type-specific markers expressed in liver, iWAT, and spleen, respectively.

We performed quantitative bottom-up proteomic analysis of labeled proteins from individual tissues and plasma using Tandem Mass Tag-based multiplexing^23^ (TurboID-TMT). Biotinylated proteins were enriched with SA beads from iWAT, liver, and spleen lysates (Figure 1E), followed by reduction, alkylation, and trypsin digestion. After TMT labeling (Figure S1E) samples were combined, fractionated to increase proteome coverage, and analyzed by mass spectrometry. Cre-negative controls were used to exclude nonspecific and endogenously biotinylated proteins. Receiver operating characteristic (ROC)-based analysis was applied as previously described^11^ to define specific thresholds identifying ER-localized proteins with fidelity (Figures S1F-S1G, see Supplementary Methods for details). As expected, proteins enriched in Cre+ samples showed Gene Ontology associations with subcellular localization to the secretory pathway (ER, extracellular, and Golgi). Further many of the proteins were predicted by SignalP^24^ to contain signal peptides (Figures 1F, S1H, S1I). Finally, the secretory proteomes from adipocytes, hepatocytes, and B cells were clearly separated by principal component analysis (PCA) (Figure S1J) and included numerous cell type-enriched proteins annotated in the Human Protein Atlas (Figure 1G, Table S1). Together, these studies validated the use of the genetically encoded TurboID^KDEL^ mouse strain combined with our proteomic pipeline as a robust and versatile platform for characterizing the secreted proteome *in vivo* as well as ER-resident, and ER-transitory proteins such as glycoproteins and membrane receptors.

### *In vivo* models for profiling secretory pathway proteomes under negative-energy balance

We next used this approach to characterize the secretory proteome in adipose tissue and liver under two conditions of negative energy balance: a) a 48-hour fast and b) lipopolysaccharide (LPS) treatment (1 mg/kg, intraperitoneally once daily for 48 hours) to induce inflammation-associated anorexia.^7^ Both perturbations were compared to *ad libitum*-fed, saline-injected controls. Mice were concurrently provided with biotin-supplemented drinking water (0.25 mg/mL) for 2 days, enabling TurboID-mediated biotinylation *in vivo* (Figure 2A).

**Figure 2.**
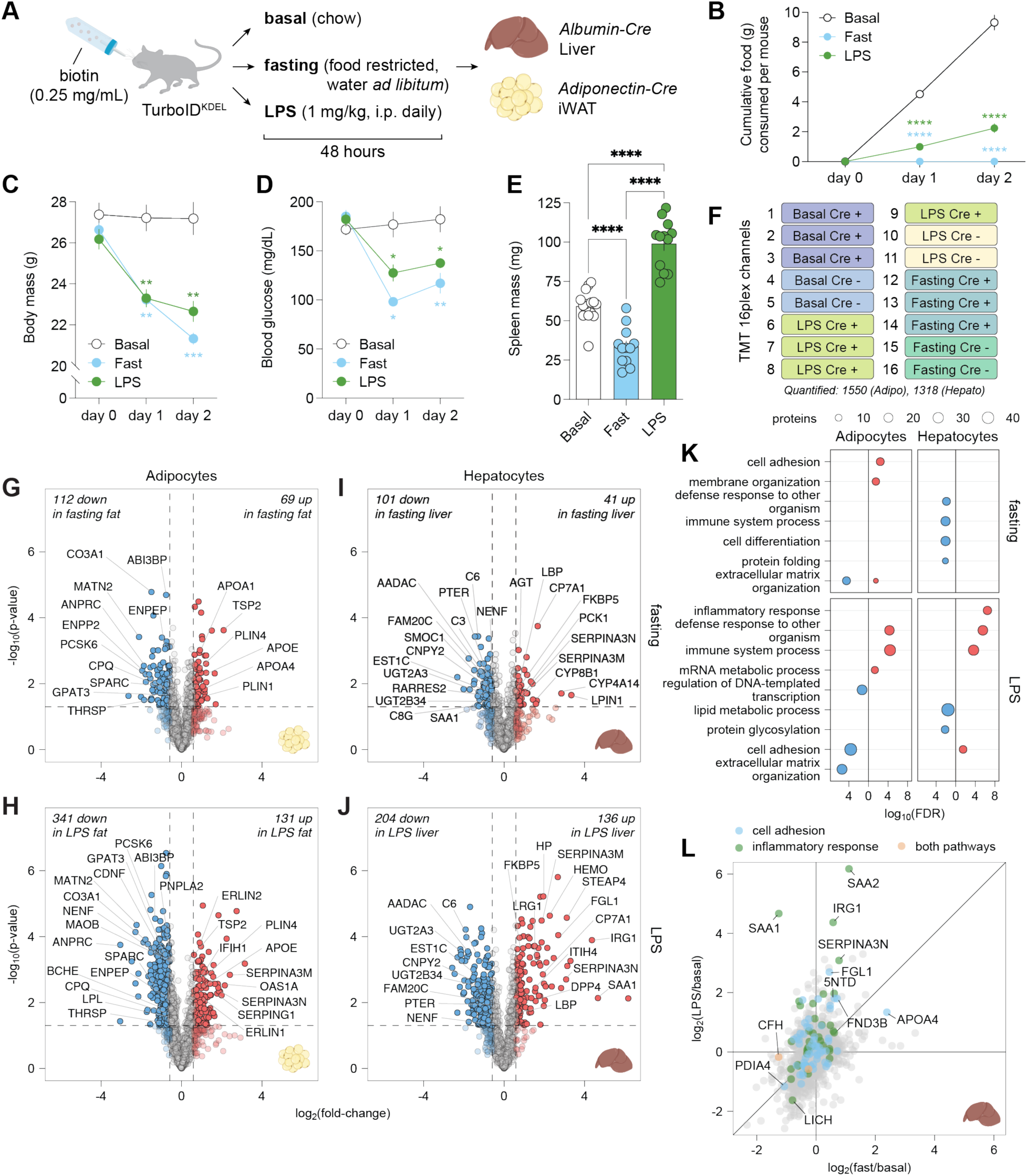
Proteomic response of hepatocytes and adipocytes to negative energy balance. (A) Experimental design to induce negative energy balance in mice, including 48 hours of LPS treatment or fasting in two tissue-specific driver lines, Adiponectin-Cre (adipose tissue) and Albumin-Cre (hepatocytes). (B) Cumulative food consumption in LPS and fasting conditions over 48 hours. LPS and fasting were each compared to the basal condition by two-way ANOVA with Tukey’s multiple comparisons test (****, p < 0.0001). Error bars show mean ± SEM; n = 6-8 cages per condition. Mice include Cre+, Cre-, and wild-type genotypes. (C) Body mass change over 48 hours of fasting or LPS. Conditions compared to basal by two-way ANOVA with Tukey’s multiple comparisons test (ns, p > 0.05, not shown; ** p < 0.01; *** p < 0.001). Error bars show mean ± SEM; n = 6-8 mice per condition. Mice include Cre+, Cre-, and wild-type genotypes. (D) Change in blood glucose over 48 hours of fasting or LPS. Conditions compared to basal by two-way ANOVA with Tukey’s multiple comparisons test (ns, p > 0.05, not shown; *, p < 0.05; **, p < 0.01). Error bars show mean ± SEM; n = 6-8 mice per condition, including Cre+, Cre-, and wild-type genotypes. (E) Spleen mass following 48 hours of fasting or LPS. Conditions compared by one-way ANOVA with Tukey’s multiple comparisons test (****, p < 0.0001). Error bars show mean ± SEM; n ≥ 12 mice per condition. (F) Representative TMT channel assignment for a 16-plex LC-MS/MS/MS experiment comparing three conditions (basal, LPS, and fasting) in mice with and without a cell-type-specific Cre driver. The number of proteins passing the ROC cutoff is also shown. (**G-H**) Volcano plots showing log_2_ fold changes in protein expression in inguinal white adipose tissue (iWAT) samples from fasted or LPS-treated Adipo-TurboID^KDEL^ mice relative to basal. Dashed lines represent cutoffs of p-value < 0.05 and FC > 1.5. (**I-J**) Volcano plots showing log_2_ fold changes in protein expression in liver tissue samples from fasted or LPS-treated Albumin-TurboID^KDEL^ relative to basal. Dashed lines represent cutoffs of p-value < 0.05 and fold-change > 1.5. (K) Pathway analysis of protein expression changes. Dot plots show GO terms with < 0.05 FDR when using Fisher’s exact test against an experiment-specific background list. (L) Scatter plot showing protein expression changes in liver tissue samples from fasted (x-axis) or LPS-treated (y-axis) Albumin-TurboID^KDEL^ mice. Select pathways are labeled based on Gene Ontology annotations.

Cumulative food intake was significantly reduced in LPS-treated mice compared to controls (Figure 2B). Both interventions led to significant decreases in body mass and blood glucose (Figures 2C, 2D). While both sets of conditions are associated with negative energy balance, we also found that LPS induced splenomegaly indicative of selective activation of immune responses, which was not observed in fasted mice (Figure 2E).

Following each perturbation, biotinylated proteins were enriched from iWAT and liver from Adipo-TurboID^KDEL^ and Albumin-TurboID^KDEL^ mice, respectively, and quantified by TurboID-TMT proteomics (Figure 2F, Tables S1, S2). MS identified 2,460 proteins in iWAT lysates, of which 1,550 passed the ROC-based enrichment threshold. Liver samples were processed similarly in a separate TurboID-TMT 16-plex experiment, with 2,608 proteins identified in liver lysates, of which 1,318 met the ROC-based enrichment threshold. As expected, many enriched proteins contained predicted signal peptides by SignalP (376 in iWAT and 343 in liver; Table S2).

### Fasting and LPS trigger distinct remodeling of the secretory proteome in adipocytes

Fasting had highly significant effects on the white adipocyte proteome, with 181 differentially expressed proteins (p < 0.05; fold change, FC >1.5) (Figure 2G). Fasting reduced abundance of ECM components, including ABI3BP (-0.8 log_2_FC), MATN2 (-2.1 log_2_FC), SPARC (-1.2 log_2_FC), and CO3A1 (-1.5 log_2_FC) suggesting that it induced extensive extracellular matrix (ECM) remodeling. Concomitantly, proteins involved in lipid synthesis and storage (GPAT3, -0.6 log_2_FC; THRSP, -1.5 log_2_FC) and adipose tissue expansion (ENPP2, -2.0 log_2_FC) were downregulated. In contrast, fasting upregulated lipid metabolism and transport proteins, including apolipoproteins (APOE, 0.7 log_2_FC; APOA1, 0.9 log_2_FC; APOA4, 1.2 log_2_FC) and perilipins (PLIN1, 1.2 log_2_FC; PLIN4, 0.9 log_2_FC), consistent with lipid mobilization from adipose stores. In addition, several proteases and aminopeptidases (including CPQ, -0.7 log_2_FC; ENPEP, -0.8 log_2_FC; and PCSK6, -0.8 log_2_FC) were downregulated; while fasting is known to induce profound metabolic reprogramming in adipose tissue,^25^ the reduced expression of these enzymes may suggest diminished peptide processing and proteolytic activity under nutrient deprivation.

Acute LPS treatment induced even more extensive changes of the adipocyte secretory proteome (Figure 2H), with 341 proteins significantly downregulated and 131 upregulated (p < 0.05; FC > 1.5). Consistent with the negative energy balance observed in both conditions, LPS-treated adipocytes also showed downregulation of proteins involved in lipid metabolism, including LPL (-0.6 log_2_FC), PNPLA2 (-0.6 log_2_FC), GPAT3 (-0.8 log_2_FC), and THRSP (-1.0 log_2_FC), reduced expression of ECM components,^26^ and upregulation of the adipogenesis inhibitor TSP2 (0.6 log_2_FC).

LPS also induced several markers of inflammation in adipose tissue including interferon-stimulated genes (IFIH1, 0.9 log_2_FC; OAS1A, 2.2 log_2_FC) and negative regulators of inflammation, such as serine protease inhibitor family members (SERPINA3N, 1.9 log_2_FC; SERPINA3M, 2.4 log_2_FC; SERPING1, 1.8 log_2_FC). We also observed increased ER Lipid Raft-Associated Proteins 1 and 2 (ERLIN1 and 2, 1.0 and 0.9 log_2_FC, respectively), suggesting activation of mechanisms that mitigate ER stress which potentially results from the marked induction of many proteins.^27,28^ Like fasting, LPS treatment led to downregulation of proteases, including CPQ (-0.7 log_2_FC), ENPEP (-0.6 log_2_FC), and PCSK6 (-1.5 log_2_FC). However, unlike fasting, LPS also induced a significant decrease in secretion of neurotrophic factors (CDNF, -1.3 log_2_FC; NENF, -1.1 log_2_FC) and proteins involved in neurotransmitter degradation (MAOB, -0.6 log_2_FC; BCHE, -0.9 log_2_FC).

### Fasting and LPS trigger distinct remodeling of the hepatocyte secretory proteome

Fasting induced differential expression of 142 hepatocyte secretory pathway proteins (p < 0.05; FC >1.5) (Figure 2I; Table S3). Multiple complement components (C3, -0.8 log_2_FC; C6, -1.3 log_2_FC; C8G, - 1.3 log_2_FC) and the acute-phase hepatokine SAA1 (-1.3 log_2_FC) were downregulated in line with suppression of immune responses during fasting^29,30^ and the central role of hepatocytes in complement activation and acute phase responses. Proteins involved in lipid catabolism and energy conservation were also decreased, including AADAC (-0.9 log_2_FC), EST1C (-0.7 log_2_FC), and PTER (-0.8 log_2_FC).^31^ Other notable decreases include neurotrophic and growth-related factors (NENF, -0.6 log_2_FC; CNPY2, -0.8 log_2_FC), the glucose-responsive hepatokine SMOC1 (-0.7 log_2_FC), and the secretory pathway kinase FAM20C (-0.9 log_2_FC), which regulates ECM remodeling. Consistent with the role of the liver in phase II detoxification via glucuronidation, ER-localized glucuronosyltransferases (including UGT2A3, -0.8 log_2_FC; UGT2B34, -0.8 log_2_FC) were suppressed. In contrast, fasting proteins involved in gluconeogenesis were induced together with enzymes involved in bile acid biosynthesis and fatty acid metabolism (CP7A1, 1.7 log_2_FC; PCK1, 1.3 log_2_FC; CYP8B1, 1.6 log_2_FC; CYP4A14, 2.8 log_2_FC; and LPIN1, 3.3 log_2_FC). Overall, these data were consistent with metabolic reprogramming toward fuel production which differed from the response in adipocytes (Table S3).

LPS led to broader changes of the liver secretory pathway proteome compared to fasting (Figure 2J), inducing differential expression of 340 proteins (p < 0.05; FC >1.5). This included upregulation of acute-phase proteins (LBP,^32^ SAA1,^33^ HEMO, HP; log_2_FCs of 0.6, 4.7, 2.3, and 2.0, respectively), anti-inflammatory modulators (SERPINA3M, SERPINA3N, FGL1; log_2_FCs of 1.9, 3.1, and 2.7, respectively), and integrators of inflammatory and metabolic signaling (DPP4, STEAP4; log_2_FCs of 1.6 and 3.1, respectively). We also noted upregulation of leucine-rich α2-glucoportein 1 (LRG1, log_2_FC 1.5),^34,35^ a factor implicated in hepatic steatosis and insulin resistance.

### Shared and tissue-specific responses to fasting and LPS in secretory proteomes

To capture shared and distinct responses across cell types and negative energy balance models, we performed Gene Ontology (GO) pathway analysis of “biological processes” enriched in adipocyte and hepatocyte secretory pathway proteomes following fasting or LPS (Figures 2K, 2L, S2A-S2F). As expected, LPS promoted inflammatory and defense response pathways in both cell types (Figures 2K, 2L, S2A). In contrast, immune-related pathways were downregulated in fasted hepatocytes.^36,37^

Proteins regulating cell adhesion and ECM organization were the most enriched processes in adipocyte-derived secretory proteomes, responding distinctly between fasting and inflammatory stress (Figures 2K, S2B). Key quality control processes mediated in the ER, including protein folding and protein glycosylation, were also significantly enriched in both hepatocytes and adipocytes (Figures 2K, S2C, S2D), while lipid metabolic processes were downregulated in LPS-stimulated hepatocytes (Figures 2K, 2L).

Comparison of liver and adipose secretory pathway proteins highlighted the shared and tissue-specific responses to fasting and LPS (Figures S2F-S2H). LPS induced inflammatory regulators (SERPINA3N, SERPINA3M, SAMP) and downregulated lipid anabolic enzymes, including the major lipase CES1D in both experiments, while many of the acute phase response proteins were only detected in liver samples. We validated these findings by performing Western blot analysis of independent wild-type mouse cohorts treated with biotin during fasting or LPS treatment; CES1D and C6, both downregulated in LPS and fasting conditions in the liver by MS, were consistently reduced by Western blotting (Figures S2I, S2J).

### Changes in circulating hepatokines and adipokines in response to negative energy balance

Plasma proteins were collected from basal, LPS-injected, and fasted Albumin-TurboID^KDEL^ and Adipo-TurboID^KDEL^ mice enriched on SA beads, TMT-labeled, and analyzed by mass spectrometry to define cell-type-specific circulating secretory proteins (Figures 3A-3D, S3A-S3C), hereafter referred to as the ‘hepato-plasma proteome’ and ‘adipo-plasma proteome’, respectively. Consistent with selective capture of circulating proteins, plasma datasets were significantly enriched for proteins predicted to have a signal peptide (>70%; Figure 3B, Table S1).

**Figure 3.**
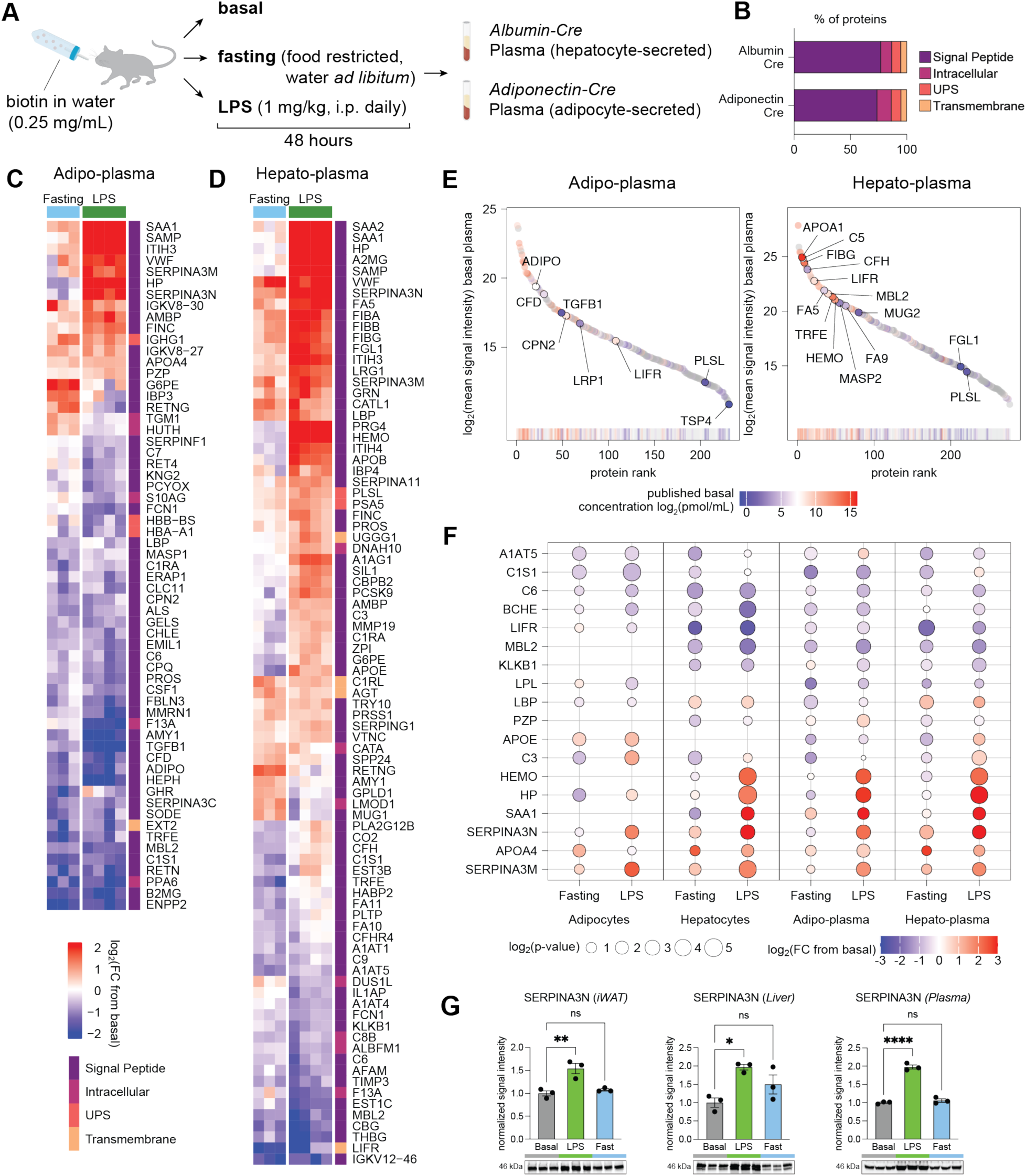
Hepato-plasma and adipo-plasma proteomic responses to negative energy balance. (A) Experimental design to capture circulating adipokines and hepatokines in response to fasting or LPS in TurboID mice. (B) Bar plot of proteins quantified in Albumin-TurboID^KDEL^ and Adipo-TurboID^KDEL^, with protein class labeled based on SignalP and OutCyte predictions. (C) Heatmap of differentially expressed proteins in adipo-plasma (p < 0.05, FC > 1.5 in either of the negative-energy-balance conditions compared to basal). Proteins are hierarchically clustered based on FC. (D) Heatmap of differentially expressed proteins in hepato-plasma (p < 0.05, FC > 1.5 in either of the negative-energy-balance conditions compared to basal). Proteins are hierarchically clustered based on FC. (E) Rank plot of mean signal intensity values of proteins quantified by TurboID-TMT in basal adipo- and hepato-plasma colored by known plasma concentrations. (F) Dot plot of log_2_FC and p-values of selected proteins detected in adipocytes, hepatocytes, and plasma across negative energy balance conditions. (G) Validation by Western blot of SERPINA3N upregulation in response to LPS in bulk iWAT, liver, and plasma. Western blot band intensity was normalized to the total intensity of the corresponding lane in a stain-free gel image. Comparison performed by one-way ANOVA with Dunnett’s multiple comparisons test (ns, p>0.05; *, p<0.05; **, p < 0.01; ****, p < 0.0001).

To evaluate the dynamic range of plasma proteins detected, we compared basal plasma reporter ion intensities for 279 proteins that were found to intersect with a published atlas of mouse plasma proteome with reported absolute concentrations^38^ (Figures 3E, S3D, S3E). Proteins with the highest concentrations included apolipoprotein A-I (APOA1, 47.5 nmol/mL) and fibrinogen gamma chain (FIBG, 9.3 nmol/mL), whereas detected proteins with the lowest concentrations were plastin-2 (PLSL, 1.4 pmol/mL), fibrinogen-like protein 1 (FGL1, 0.8 pmol/mL), and thrombospondin 4 (TSP4, 0.7 pmol/mL), suggesting sub-nanomolar sensitivity. Albumin was also one of the top quantified proteins, which was filtered out as a common MS contaminant^39^ in our analysis pipeline, and has a basal concentration of 137.3 nmol/mL. While relatively non-abundant in total plasma compared to liver, several proteins expressed by adipocytes, including LRP1 and CFD (7.8 and 67.2 pmol/mL, respectively), were detected in the adipo-plasma proteome (Figures 3E, S3D), further validating the enrichment strategy for detecting secreted factors from their tissues of origin even at low abundance. Hepatocyte-enriched proteins FA5, FA9, MASP2, and CFH were similarly overrepresented in the hepato-plasma proteome by TurboID^KDEL^ relative to absolute concentrations from total plasma (Figures 3E, S3E).

A subset of biotinylated proteins we detected in liver and/or iWAT tissues were also found in the plasma-derived samples (Figures 3F, S3F-S3J). The intersect and concordance between tissue and plasma proteomes were higher in the Albumin-TurboID^KDEL^ (Pearson’s R = 0.74 for LPS, R = 0.68 for fasting; Figures S3I-S3J) compared to the Adipo-TurboID^KDEL^ mice (R = 0.51, R = 0.54; Figures S3G-S3H), likely reflecting differences in protein secretion kinetics and clearance rates between liver and adipose tissues, as well as potential contribution of other fat depots to the circulating adipo-proteome.

Both fasting and LPS stimulation suppressed the plasma levels of several complement components, including C6 and C1S1 (Figure 3F), which are produced in both tissues. Conversely, serine protease inhibitors SERPINA3N and SERPINA3M, which target cathepsins and granzymes, were upregulated in adipose tissue and hepatocytes raising the possibility that they serve potential protective functions during inflammatory and metabolic stress (see also Figures S3G-S3J). Likewise, APOA4 induction likely reflects a protective response to negative-energy-balance given its reported roles in lipid mobilization and anti-inflammatory signaling.

Several acute-phase and inflammation-responsive proteins, including C3, HEMO, and HP showed divergent regulation between LPS stimulation and fasting, consistent with distinct state-dependent functional roles. In contrast, the acute-phase protein LBP was decreased in both states of negative-energy balance in adipose tissue and the adipo-plasma proteome but increased in liver tissue and the hepato-plasma proteome (Figure 3F), indicating cell-specific regulation. Similarly, upon fasting, G6PE, an enzyme in the pentose phosphate pathway, was downregulated in both iWAT and adipo-plasma, but upregulated in liver and hepato-plasma (Figures S3H, S3I). We also confirmed upregulation of SERPINA3N (Figure 3F) in response to LPS by immunoblotting of tissue and plasma samples from wild-type mice (Figure 3G).

Together, the observed correlations between tissue and plasma proteomes, as well as instances of divergence under fasting and LPS stimulation, highlight the complex interplay between post-translational and systemic regulatory mechanisms governing hepatokine and adipokine secretion in response to negative energy balance.

### The white adipocyte tissue secretory proteome in mice fed a HFD for 15 weeks

We next investigated how diet-induced obesity (DIO), a state of positive energy balance during weight gain, alters the adipocyte secretome. Adipo-TurboID^KDEL^ mice were placed on a high-fat diet (HFD) for 6 or 15 weeks (Figure 4A). We then characterized the ER proteome of the two different white adipose tissue (WAT) depots known to undergo hypertrophy and hyperplasia on a HFD: subcutaneous inguinal (iWAT) and visceral epididymal WAT (eWAT). We also characterized the plasma proteome in these animals at the same timepoints.

**Figure 4.**
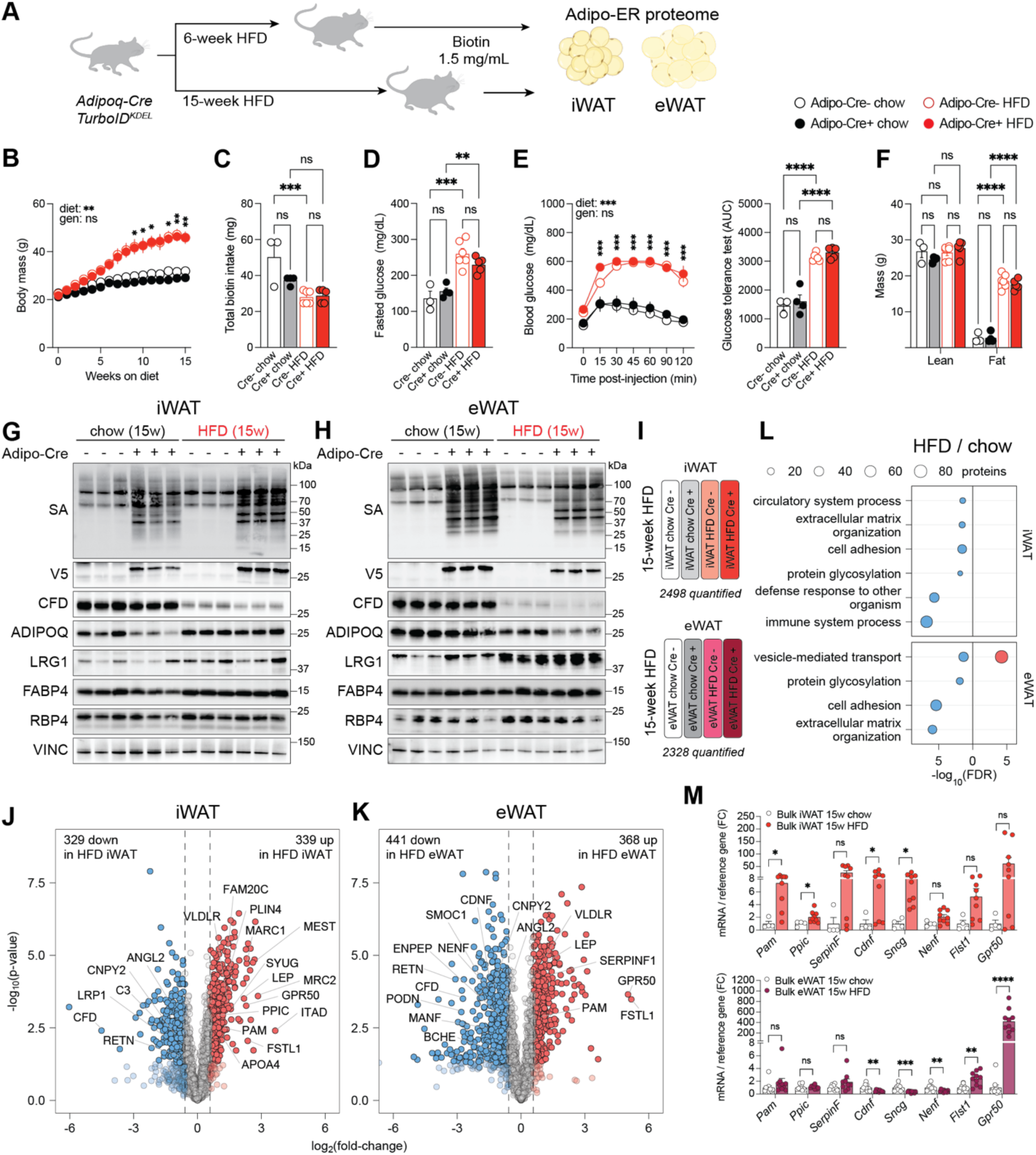
Proteomic responses of inguinal and epidydimal adipocytes in Adipo-TurboID^KDEL^ mice to advanced obesity. (A) Experimental design to capture the secretory pathway proteome in inguinal (iWAT) and epididymal white adipocytes (eWAT) of obese Adipo-TurboID^KDEL^ mice after 6 and 15 weeks of high-fat diet (HFD) feeding. (B) Body weights of mice across 15 weeks of HFD feeding. Biotin (1.5 mg/mL) was administered in drinking water for 7 days on the last week of feeding. Conditions were compared to basal by two-way ANOVA with Tukey’s multiple comparisons test (ns, p > 0.05, not shown; *, p < 0.05; **, p < 0.01). Error bars show mean ± SEM; n = 4-7 mice per condition. (C) Total biotin intake in mice after 7 days. Conditions were compared to basal ordinary one-way ANOVA with Šidák multiple comparisons test (ns, p > 0.05; *** p < 0.001). Error bars show mean ± SEM; n = 3-7 mice per condition. (D) Fasted glucose in mice fed chow or HFD for 15 weeks. Conditions were compared to basal ordinary one-way ANOVA with Šidák multiple comparisons test (ns, p > 0.05; **, p < 0.01; *** p < 0.001). Error bars show mean ± SEM; n = 3-7 mice per condition. (E) Glucose tolerance in mice fed chow or HFD for 15 weeks and area under curve (AUC). Conditions were compared to basal by two-way ANOVA with Tukey’s multiple comparisons test (ns, p > 0.05, not shown; ***, p < 0.001). Error bars show mean ± SEM; n = 3-7 mice per condition. AUC was compared using ordinary one-way ANOVA with Šidák multiple comparisons test (ns, p > 0.05; **** p < 0.0001). Error bars show mean ± SEM; n = 3-7 mice per condition. (F) Fat and lean mass of Adipo-TurboID^KDEL^ mice after 15 weeks of HFD feeding. Conditions were compared to basal by two-way ANOVA with Tukey’s multiple comparisons test (ns, p > 0.05; ****, p < 0.0001). Error bars show mean ± SEM; n = 3-7 mice per condition. (**G-H**) Western blot validation of protein biotinylation using streptavidin-HRP, tissue-specific V5 expression, and levels of known adipokines including complement factor D (CFD), adiponectin (ADIPOQ), leucine-rich glycoprotein 1 (LRG1), fatty acid-binding protein 4 (FABP4), and retinol-binding protein 4 (RBP4) in bulk iWAT (G) and eWAT (H) from Adipo-TurboID^KDEL^ mice after 15 weeks of HFD feeding. Vinculin (VINC) was used as a loading control. **(I)** Representative TMT channel assignment for a 16-plex LC-MS/MS/MS experiment comparing proteomic changes induced by 15-week HFD vs. chow in iWAT or eWAT adipocytes from Adipo-TurboID^KDEL^ Cre+ and Cre- mice (technical replicates not shown). The number of proteins passing the ROC cutoff in each experiment is also shown. (**J-K**) Volcano plots showing log_2_ fold changes in secretory pathway protein expression in iWAT (J) and eWAT (K) samples following HFD feeding for 15 weeks relative to basal. Dashed lines represent cutoffs of p-value < 0.05 and fold-change > 1.5. (**L**) Gene set enrichment analysis of biological processes differentially impacted by HFD and adipocyte source depot. Red color is enriched in upregulated genes and blue is enriched in downregulated genes (p-value < 0.05, FC > 1.5). (**M**) Gene expression of HFD-regulated proteins in iWAT and eWAT from Adipo-TurboID^KDEL^ mice after 15 weeks of HFD feeding. Conditions were compared to chow by t-test (ns, p > 0.05; *, p < 0.05; **, p < 0.01; ***, p < 0.001; ****, p < 0.0001). Error bars show mean ± SEM; n = 4-7 mice per condition.

Mice fed a 60% HFD for 6 weeks (Figure S4A) or 15 weeks (Figure 4B) showed significant weight gain compared to chow-fed littermates, independent of TurboID genotype. All mice received biotin in drinking water (1.5 mg/mL) for 7 days prior to tissue and plasma collection for MS analysis, and no differences in biotin-water intake were observed compared to Cre-negative controls (Figures 4C, S4B). As expected, mice fed HFD at both timepoints exhibited higher fasted glucose (Figures 4D, S4C), impaired glucose disposal during a glucose tolerance test (GTT) (Figures 4E, Figure S4D), and elevated fat mass (Figure 4F) compared to chow-fed controls.

We confirmed TurboID expression by V5 Western blotting and verified robust proteome biotinylation in bulk iWAT and eWAT from mice at 6 and 15 weeks of DIO (Figures 4G, 4H, S4E, S4F). We next compared the expression levels of adipocyte proteins known to change in abundance in DIO mice by performing Western blots on whole tissue extracts. Consistent with prior reports, we observed alterations in the levels of the following proteins in tissue derived from the 15-week DIO group: complement factor D (CFD or adipsin), LRG1, fatty acid binding protein 4 (FABP4), and retinol-binding protein 4 (RBP4)^34,40–42^ (Figures 4G, 4H).

We then analyzed the biotinylated proteomes from iWAT and eWAT lysates from 15-week HFD-fed mice using TurboID-TMT, enabling comparisons within each depot across two 16-plex experiments of animals on chow and HFD (Figure 4I; technical replicates not shown). Using experiment-specific ROC cutoffs, we quantified more than 2,300 unique proteins in both iWAT and eWAT samples at 15 weeks of chow vs. HFD feeding (Figure 4I; Table S1). Among these, 668 iWAT proteins and 809 eWAT proteins showed significant diet-specific regulation (p < 0.05, FC > 1.5; Figures 4J, 4K). GO term analysis revealed proteins reduced in the DIO samples were associated with glycosylation and ECM formation in both adipose depots (Figure 4L). In the 15-week HFD samples, 80 proteins associated with vesicle-mediated transport were specifically upregulated in eWAT adipocytes, while in iWAT adipocytes there was downregulation of genes associated with immune system processes not seen in eWAT samples.

More than 300 proteins were significantly elevated in each depot in DIO mice, including leptin (LEP; 2.2 log_2_FC in iWAT, 1.2 log_2_FC in eWAT), perilipin 4 (PLIN4; 1.4 log_2_FC in iWAT, 0.5 log_2_FC in eWAT), follistatin-related protein 1 (FSTL1; 2.4 log_2_FC in iWAT, 5.2 log_2_FC in eWAT), and serpin peptidase inhibitor 1 (SERPINF1; 1.3 log_2_FC in eWAT). We also detected decreased levels of CFD (-4.5 log_2_FC in iWAT, -2.7 log_2_FC in eWAT) and resistin (RETN; -0.8 log_2_FC in iWAT, -3.0 log_2_FC in eWAT), a hormone involved in insulin resistance (Figures 4J, 4K). Additional factors previously annotated as adipocyte-derived or obesity-regulated (but not characterized) were also identified including SMOC1 (-0.3 log_2_FC in iWAT, -1.9 log_2_FC in eWAT), peptidyl-glycine α-amidating monooxygenase (PAM; 1.1 log_2_FC in iWAT, 0.9 log_2_FC in eWAT), cerebral dopamine neurotrophic factor (CDNF; -0.3 log_2_FC in iWAT, -1.5 log_2_FC in eWAT), and γ-synuclein (SYUG; 1.6 log_2_FC in iWAT, 0.8 log_2_FC in eWAT). Finally, GPR50 was increased by 2.3 log_2_FC in iWAT and as much as 5.0 log_2_FC in eWAT, making it one of the most strongly induced proteins in both depots. GPR50 is an orphan GPCR with a sequence variant linked to elevated triglycerides and decreased HDL in obese individuals.^43^

Many of the proteins downregulated in eWAT at 15 weeks of HFD were also suppressed in mice treated with LPS, including neurotrophic factors (NENF, CDNF), metabolic (BCHE, ENPEP) and inflammatory (ANGPTL2) modulators, as well as ER stress regulators (CNPY2). These similarities between DIO and LPS treatment raise the possibility that chronic inflammation may contribute to the metabolic dysfunction associated with prolonged DIO.

To assess whether some of these protein-level changes were regulated at the transcriptional level, we performed qPCR analysis on bulk iWAT and eWAT tissues and found concordant diet- and depot-specific regulation of the genes encoding the following HFD regulated proteins: *Pam*, *Ppic*, *SerpinF*, *Cdnf*, *Sncg*, *Nenf*, *Flst1,* and *Gpr50* (Figure 4M).

### The white adipocyte tissue secretory proteome in mice fed a HFD for 6 weeks

To capture the response of adipocytes to a HFD, we characterized the adipocyte secretory proteome of iWAT and eWAT mice fed chow or HFD for 6 weeks (Figure S4G; technical replicates not shown). Western blot confirmed altered levels of CFD, LRG1, FABP4, and RBP4 expression in both depots (Figures S4E, S4F), although, consistent with prior reports, these changes were more pronounced at 15-week HFD.

Proteomic coverage was comparable across depots and diet duration, with over 2,000 proteins passing ROC cutoffs in each depot after 6 weeks on the diet (Table S1). Among these, we observed diet-specific regulation of 252 proteins in iWAT and 431 proteins in eWAT (p < 0.05, FC > 1.5; Figures S4H, S4I). Overall, similar to Western blot data, the magnitude of proteomic changes was lower than at 15 weeks. GO analysis revealed the genes regulating protein folding were significantly increased in iWAT (Figure S4J, S4K), suggesting that feeding mice a HFD for 6 weeks is associated with ER stress in this depot. Similarly, several ER stress-related proteins were upregulated in eWAT as well (Figure S4L), although less pronounced, suggesting depot-specific differences.

Consistent with the impaired glucose tolerance at 6 weeks of HFD, we found elevated levels of obesity-associated secreted factors, including LEP (2.9. log_2_FC in iWAT, 2.4 log_2_FC in eWAT), APOA4 (1.3 log_2_FC in iWAT), and VLDLR (1.0 log_2_FC in iWAT, 1.2 log_2_FC in eWAT), and reduced levels of CFD (-1.8 log_2_FC in iWAT, -2.5 log_2_FC in eWAT; Figures S4H, S4I). PLIN4 was also upregulated in both depots in HFD-fed mice; paradoxically this protein was also elevated in fat tissues from fasted mice, suggesting context-dependent roles of perilipin in the negative and positive energy balance states.

Lastly, a subset of proteins that was upregulated in iWAT at 6 weeks of HFD (Figure S4H) were downregulated in eWAT at 15 weeks of DIO (Figure 4L) as well as in the LPS-mediated negative-energy-balance model. These included the neurotrophic factor NENF, the neurotransmitter-modulating enzyme BCHE, and PAM, an enzyme which catalyzes amidation of the C-terminus of peptides which is necessary to extend the half-life of circulating factors, increasing their affinity towards targets of interest.^44^

### Depot-specific differences in DIO mice

Increased visceral adipose tissue is associated with comorbidities while subcutaneous adipose tissue is not. We investigated this by directly comparing proteomic differences between these two depots in mice fed a HFD for 6 and 15 weeks (Figures 5A-5C). Protein digests from different conditions and genotypes were split into four separate 16-plex MS experiments, which were TMT-labeled and analyzed by MS (Figures 5A, S5A; technical replicates not shown). This enabled a direct comparison of the iWAT and eWAT secretory pathway proteomes at 6 and 15 weeks under chow or HFD conditions (Figures 5B, 5C). Across experiments, we quantified between 976 and 2,373 unique proteins after ROC filters (Table S1). Combined, these proteomic datasets generated a comprehensive overview of the differences in the adipose secretory proteome in the response of each depot to the different diets. These data revealed clear depot-specific signatures (Figures 5B, 5C).

**Figure 5.**
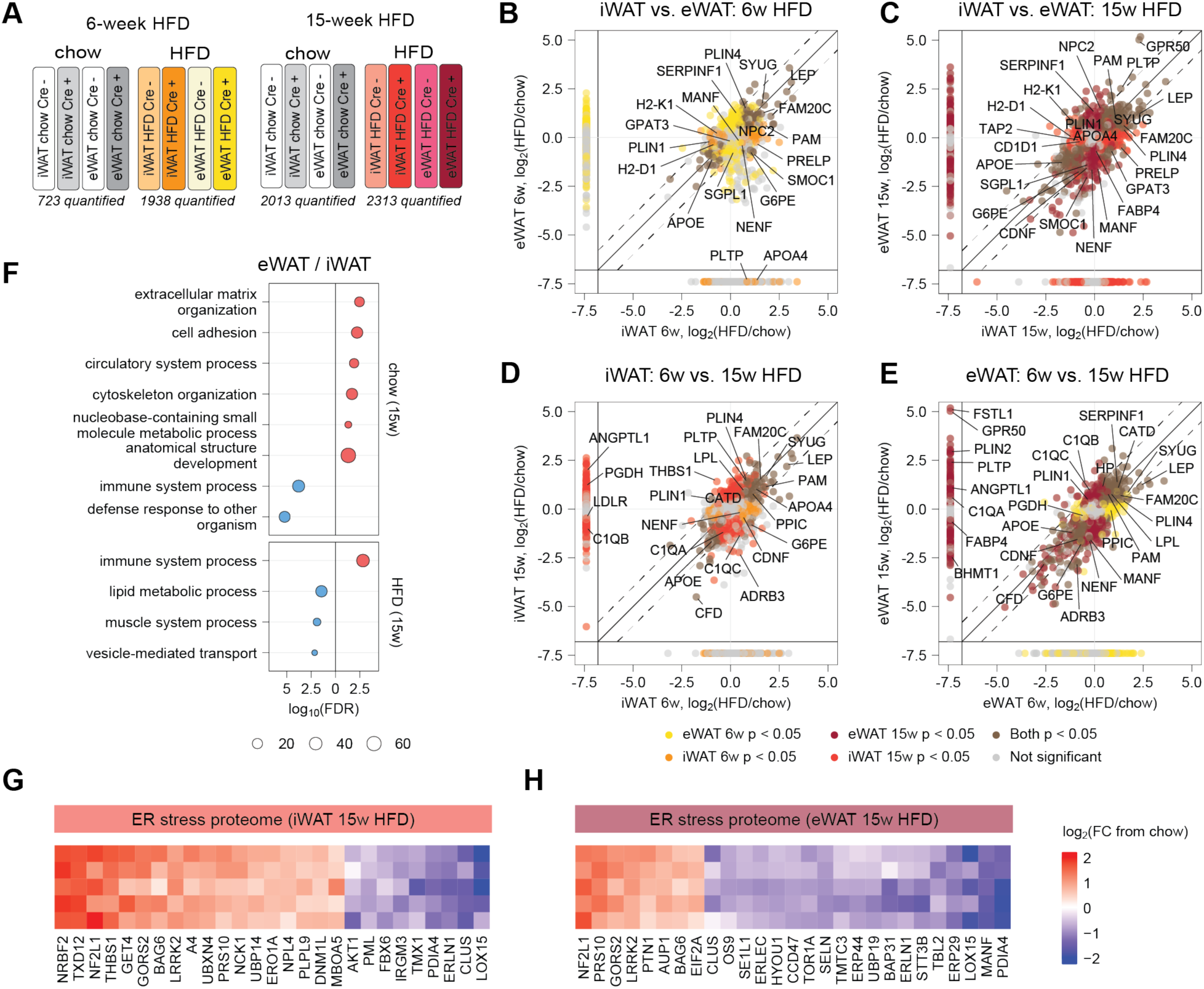
Dynamic depot-dependent reprograming of the adipocyte proteome in response to early-and late-stage DIO. **(A)** Representative TMT channel assignments for a 16-plex LC-MS/MS/MS experiment comparing iWAT and eWAT-derived adipocyte proteomes of Adipo-TurboID^KDEL^ Cre+ and Cre- mice fed chow or HFD for 6 or 15 weeks (technical replicates not shown). The number of proteins passing the ROC cutoff in each experiment is also shown. (**B-C**) Correlational analysis of inter-depot differences (inguinal vs. epididymal) in log_2_ fold changes in protein expression at 6 weeks (B) and 15 weeks (C) of HFD feeding compared to chow. Proteins quantified in only one of the fat depots are shown on margins. (**D-E**) Correlational analysis of log_2_ fold changes in protein expression following HFD feeding for 6 and 15 weeks in inguinal (D) and epididymal (E) adipocytes compared to chow. Proteins quantified for only one of the HFD timepoints are shown on margins. (**F**) Gene set enrichment analysis of biological processes differentially impacted by HFD in eWAT and iWAT adipocytes after 15-week HFD feeding compared to chow. Red color is enriched in upregulated genes and blue is enriched in downregulated genes (p-value < 0.05, FC > 1.5). (**G-H**) Heat maps showing differentially expressed proteins contributing to ER stress GO-term enrichment in iWAT (G) and eWAT (H) samples following HFD feeding for 15 weeks. Data is presented as log_2_(fold-change) compared to chow condition.

In the eWAT proteome, there was specific upregulation of the cholesterol-trafficking protein NPC2 (log_2_FC of 0.6, 0.3, 0.7, and 1.4 in 6w iWAT, 6w eWAT, 15w iWAT, and 15w eWAT, respectively) and anti-angiogenic factor SERPINF1 (log_2_FC of 0.4, 1.2, and 1.3 in 6w iWAT, 6w eWAT, and 15w eWAT, respectively), accompanied by downregulation of neurotrophic factor CDNF (-0.3 and -1.5 log_2_FC in 15w iWAT and eWAT, respectively), SMOC1 (-0.3 and -1.9 log_2_FC in 15w iWAT and eWAT of DIO), and sphingosine-1-phosphate lyase SGPL1 at 15 weeks of DIO (-0.3 and -1.3 log_2_FC in iWAT and eWAT). In contrast, the following proteins were specifically upregulated by DIO in iWAT, but not eWAT (Figures 5B, 5C): APOA4 (1.3 and 1.2 log_2_FC at 6 and 15 weeks, respectively), the insulin receptor-stabilizing factor CAVIN2^45^ (0.9 and 0.7 log_2_FC at 6 and 15 weeks, respectively), and the multifunctional ECM protein PRELP (1.3 log_2_FC at both 6 and 15 weeks in iWAT). In the secretory proteome of iWAT there was also an apparent coordinated suppression of antigen-presentation machinery in response to DIO not seen in the eWAT secretory proteome. This included downregulation of the glycolipid-presenting protein CD1D1 (-0.3 and -0.6 log_2_FC at 6 and 15 weeks of DIO) and multiple MHC class I components – peptide-loading complex (TAP2, TPSN; -0.7 and -0.9 log_2_FC in 15w iWAT, respectively) and MHC class I molecules (H2-D1, H2-K1; -1.1 and -1.1 log_2_FC in 15w iWAT, respectively).

We next compared proteomic changes between 6 and 15 weeks of DIO from separate MS experiments (Figures 5D, 5E). Many proteins were similarly altered by diet independent of duration, including CFD, ADRB3, and LEP. However, a subset of the proteome was only altered at one timepoint and not the other with a larger number of proteins showing a greater increase at 15 weeks including the lysosomal protease CATD (0.7 and 1.0 log_2_FC in 15w iWAT and eWAT, respectively), acute-phase protein HP (1.0 log_2_FC in 15w eWAT), and complement components (C1QA, C1QB, C1QC; 0.7, 1.3 and 0.9 log_2_FC in 15w eWAT, respectively). Many of the proteins that were upregulated at 15 weeks were not detected at 6 weeks suggesting that they may play a role in the adaptation to increasing adiposity. These included the anti-angiogenic factor ANGPTL1, the pro-inflammatory cytokine FSTL1, PLIN2, and GRP50. Proteins quantified in only a single dataset are shown on the x- and y-axes (Figures 5B-5E), although their absence in the 6-week experiment prevented direct temporal comparison.

Conversely, we identified a subset of proteins specifically regulated at the 6-week timepoint or showing divergent regulation across timepoints, including an increase in G6PE at 6 weeks in iWAT, while it was significantly decreased at 15 weeks of DIO in both depots (Figures 5D, 5E). The following proteins were also elevated at 6 weeks of DIO vs. 15-week samples including neurotrophic factors CDNF, MANF, and NENF which were increased in iWAT adipocytes relative to chow but decreased in eWAT adipocytes (Figure S5B). Moreover, by 15 weeks of DIO, these factors were significantly decreased in eWAT adipocytes but not in iWAT, suggesting a depot-specific response to HFD. GO analysis further revealed that in eWAT adipocytes HFD elicited a stronger increase in proteins linked with immune processes compared to iWAT, while under chow conditions, the opposite pattern was observed (Figures 5F, S5C, S5D). A similar induction of proteins associated with an immune response was also observed in iWAT after the LPS-treatment even though the two states are quite distinct. Finally, multiple proteins associated with ER stress were significantly altered in both depots in late-stage obesity following 15 weeks of HFD feeding (Figures 5G, 5H).

### Plasma proteins secreted by adipose tissue in early and advanced obesity

We next performed TurboID-TMT analysis of plasma proteins secreted by adipose tissue of DIO obese mice (Figure 6A) after 6 and 15 weeks of HFD feeding (Figure 6B; technical replicates not shown). As expected, plasma samples were significantly enriched for SignalP-containing proteins, confirming selective capture of secreted proteins, including LEP, CFD, LRG1, PLTP, as well as unconventionally secreted proteins (UPS) such as SYUG and PPIC (Figures 6C, S6A).

**Figure 6.**
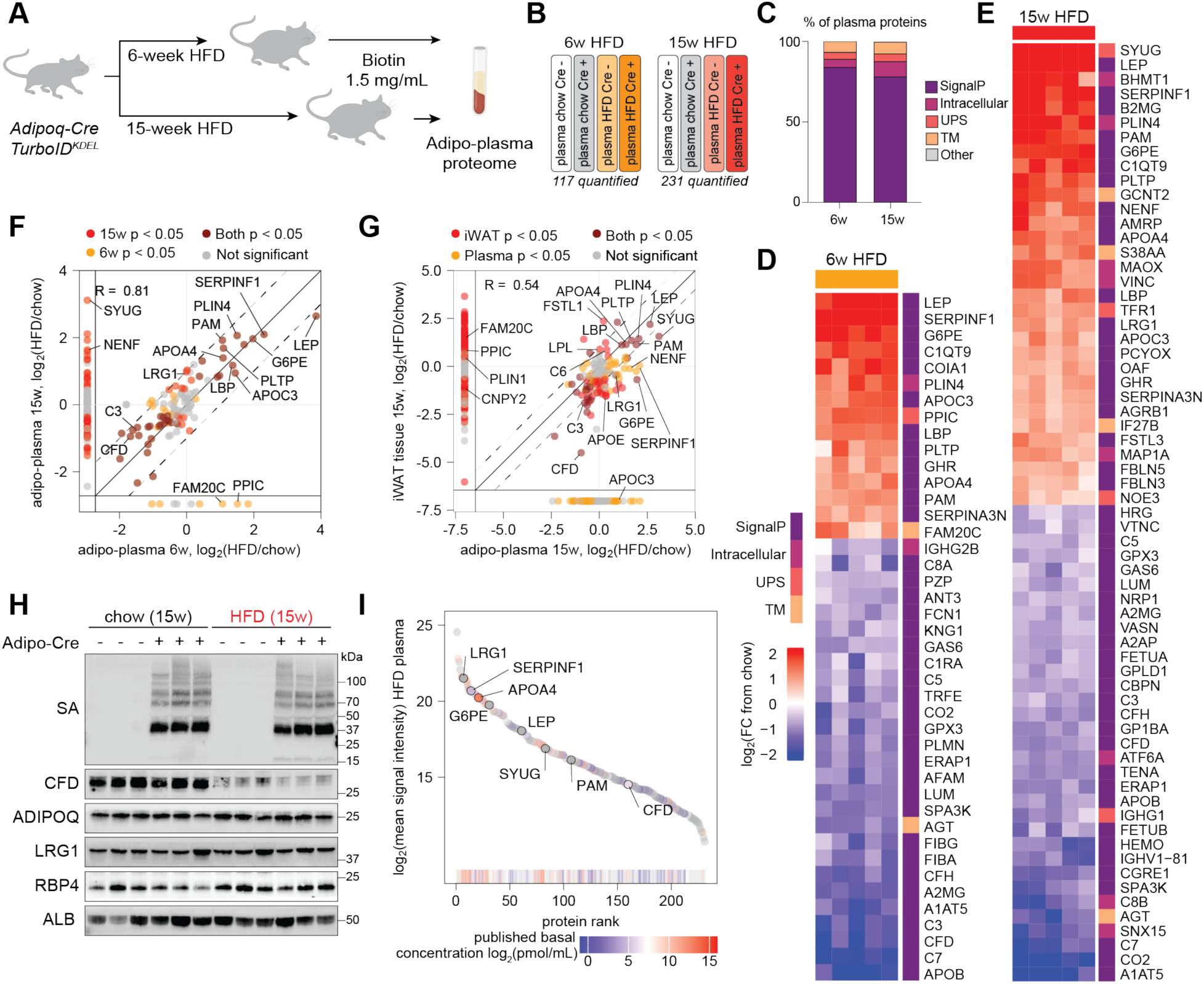
Adipo-plasma proteome in early and advanced DIO. **(A)** Experimental design for the discovery of circulating adipokines in obese Adipo-TurboID^KDEL^ mice after 6 and 15 weeks of HFD feeding; n = 5 mice per condition. (**B**) Representative TMT channel assignments for a 16-plex LC-MS/MS/MS experiment comparing adipo-plasma proteomes of Adipo-TurboID^KDEL^ Cre+ and Cre- mice fed chow or HFD for 6 or 15 weeks (technical replicates not shown). The number of proteins passing the enrichment cutoff in each experiment is also shown. (**C**) Bar plot of proteins quantified in Adipo-TurboID^KDEL^ plasma proteome at 6 and 15 weeks of DIO, with protein class labeled based on SignalP and OutCyte predictions. **(D-E)** Heatmaps showing Adipo-TurboID^KDEL^ plasma proteome affected by HFD feeding in early (D) and late (E) stage of obesity. Data is presented as log_2_(fold-change) compared to chow condition. (**F**) Scatterplot of log_2_ fold changes in adipo-plasma protein expression following HFD feeding for 6 and 15 weeks compared to chow. Proteins quantified for only one of the HFD timepoints are shown on margins. (**G**) Scatterplot of log_2_ fold changes in protein expression in iWAT vs. plasma proteomes of Adipo-TurboID^KDEL^ mice following HFD feeding for 15 weeks compared to chow. Proteins quantified for only one of the conditions are shown on margins. (**H**) Western blot validation of protein biotinylation using streptavidin-HRP and levels of known adipokines including CFD, ADIPOQ, LRG1, and RBP4 in bulk plasma samples from Adipo-TurboID^KDEL^ mice after 15 weeks of HFD feeding. Albumin (ALB) was used as a loading control. (**I**) Rank plot of mean signal intensity values of proteins quantified by TurboID-TMT in Adipo-TurboID^KDEL^ plasma proteome following HFD feeding for 15 weeks colored by known basal plasma concentrations.

We identified 117 proteins from the adipo-plasma proteomes of Adipo-TurboID^KDEL^ mice following 6 weeks and 213 proteins after 15 weeks of high fat diet (Table S1). Of these, 42 plasma proteins were significantly changed at 6 weeks of DIO (15 up and 27 down), and 65 were altered at 15 weeks (32 up and 33 down in HFD vs. chow) (p < 0.05, FC > 1.5; Figures 6D-6F). These included canonical adipose-derived hormones such as LEP, as well as other adipokines sensitive to nutritional state, including CFD and its cleavage product C3, phospholipid transfer protein PLTP, PLIN4, LBP, SERPINF1, and APOA4. Comparison of adipo-plasma proteomes between early- and late-stage obesity (Figure 6F) revealed that a large subgroup of known adipokines were increased in the samples from mice fed a HFD for 15 weeks including LEP, SERPINF1, CFD, and G6PE. However, a smaller number of proteins were detected only at one stage. These included proteins specifically detected at 6 weeks of HFD feeding such as FAM20C, and some mediators such as SYUG and BHMT1 which were only increased at 15 weeks of DIO and were not detected in the earlier timepoint (Fig. 6F, see also Table S1).

Consistent with our previous work, we confirmed LRG1 as an obesity-induced adipokine^34^ (Figure 6E). Moreover, SYUG, like LEP, was one of the most upregulated proteins in adipocyte-derived plasma in obese mice at 15 weeks (Figure 6E), consistent with its upregulation in both iWAT and eWAT at this timepoint (Figures 6G, S6B). We also detected increased levels of G6PE in plasma following HFD feeding, which was distinct from its decreased cellular levels in iWAT in obesity and fasting (Figure 6G), suggesting post-translational regulation of this circulating factor.

We validated biotinylation of circulating proteins in plasma, as well as diet-induced regulation of known adipokines by Western blot analysis of plasma proteins from 6- and 15-week chow and HFD fed mice, and, as expected, saw a pronounced impact of HFD on CFD levels in the chronic DIO group (15 weeks; Figure 6H) compared to 6-week fed animals (Figure S6C), which did not show a pronounced difference. Globally, analysis of the dynamic range based on signal intensity and protein rank suggested we were able to cover a wide range of abundance, including hormones and soluble factors, known to circulate in much lower concentrations than more abundant plasma proteins such as apolipoproteins, CFD or LRG1 (Figures 6I, S6D).

Notably, our experimental design enabled direct comparison of secretory pathway changes across multiple negative and positive energy-balance models (Figures S6E-S6G). This revealed both concordant effects on proteins such as G6PE, SERPINA3N, and CFD, as well as discordant regulation of proteins including GHR and B2MG across fasting, LPS, and DIO conditions.

### Secreted factors impacted by energy balance exhibit human disease associations

To investigate the potential clinical relevance of adipocyte-derived secreted proteins, we next asked which factors altered by fasting, LPS, and obesity are associated with human phenotypes in UK Biobank (UKBB), the largest plasma proteome-phenome atlas available to date. This database includes information from 53,026 individuals with Olink-based quantification of plasma proteins across 406 prevalent diseases, 660 incident diseases, and 986 health-related traits.^46^ This resource provides statistically powerful associations between basal plasma levels of a given circulating protein and prevalence of specific clinical traits and disease outcomes in human subjects.

We identified 65 proteins that were significantly changed in the adipo- or hepato-plasma proteome under low- or high-energy balance and that were associated with endocrine, circulatory, and/or infectious diseases (Figures 7A-7E, Table S4). We also found significant associations in the UKBB cohorts between obesity-regulated proteins and susceptibility to a range of infectious diseases, including LIFR, C7, and SERPINF1 (Figures 7B, 7C, S7A-S7C). Further, a subset of circulating proteins whose levels were altered by changes in energy balance exhibited strong associations and elevated odds ratios (OR) of circulatory and endocrine/metabolic diseases, including LEP, SYUG, LRG1, SERPINF1, APOA4, and BCHE (Figures 7D, 7E, S7D-S7H). Notably, T2D was among the strongest associations by adjusted p-values for SYUG (p = 3.66 x 10^-31^, OR of 1.64), SERPINF1 (p = 3.47 x 10^-40^, OR of 4.5), and APOA4 (p = 1.23 x 10^-76^, OR of 3.23) (Figures 7E, S7D, S7F, S7G), while, as expected, LEP was associated with multiple outcomes linked with obesity such as hypertension (p = 1.84 x 10^-47^, OR of 1.4), metabolic disorder (p = 6.9 x 10^-39^, OR of 1.5), and several adverse cardiometabolic outcomes.

**Figure 7.**
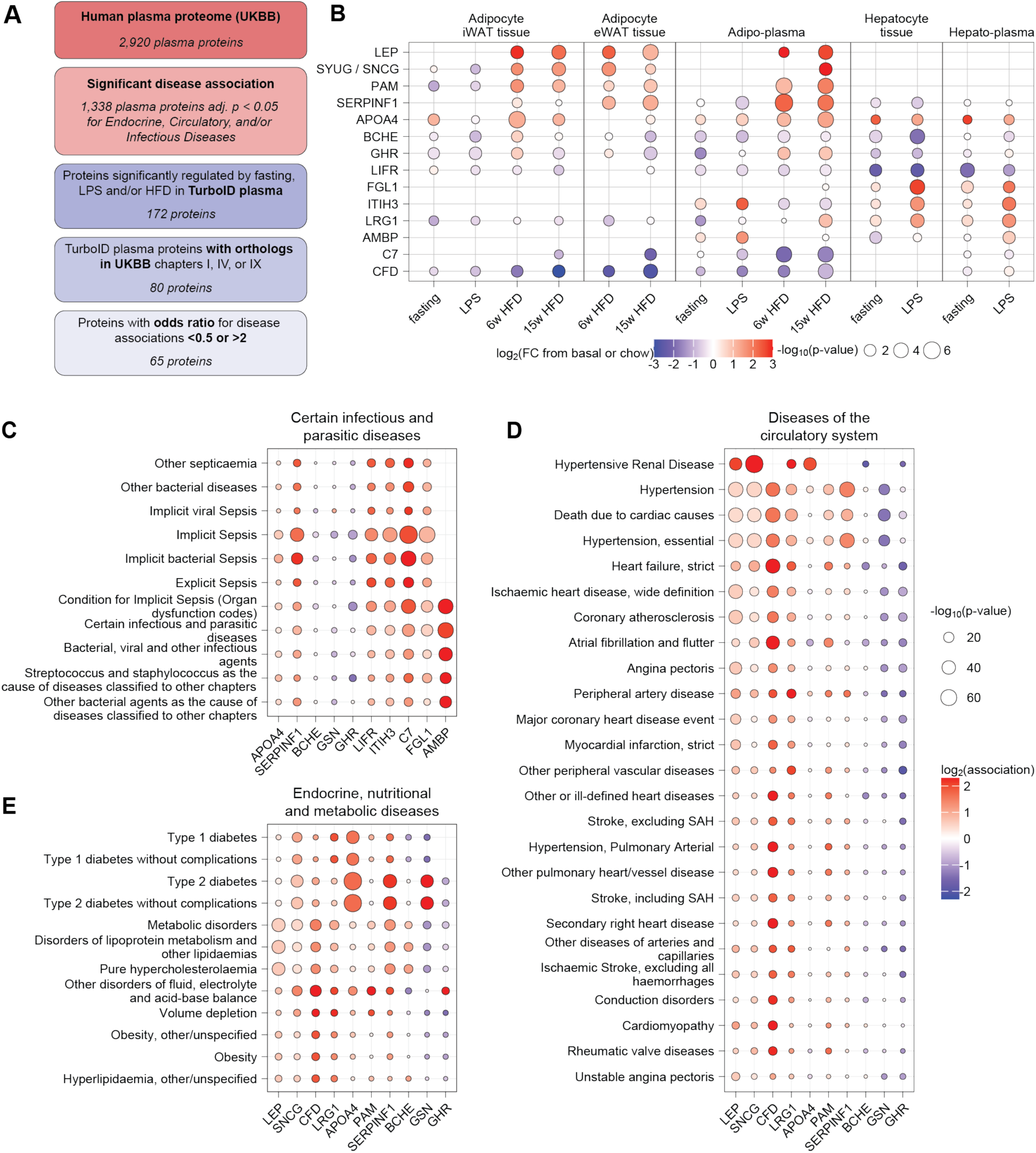
Energy balance-dependent circulating proteins are associated with human health and disease. (A) Schematic showing selection of TurboID plasma proteins with orthologs in relevant disease-associated human plasma proteins identified in a UKBB cohort of 53,026 individuals. (B) Dot plot of log_2_FC and p-values of selected proteins detected in adipocytes, hepatocytes, and plasma across negative and positive energy balance conditions. Data are shown relative to own control group (e.g. fasting vs. basal, LPS vs. basal, HFD vs. chow). (C) Infectious disease associations for top regulated hepatokines and adipokines, based on basal plasma levels of these protein and clinical outcomes in UKBB cohort of 53,026 individuals. (**D-E**) Disease associations for top regulated adipokines, based on basal plasma levels of these protein and clinical outcomes in UKBB cohort of 53,026 individuals.

Among these associations, we were particularly intrigued by human γ-synuclein (SYUG), which in our experimental data is an adipocyte-enriched protein, reduced during fasting and after LPS treatment in inguinal fat cells and increased in obesity in both fat depots and in the plasma (Figure 7B). Interestingly, UK Biobank individuals with elevated circulating levels of SYUG exhibited higher odds of several metabolic disorders including T2D, hypercholesterolemia (p = 6.64 x 10^-14^, OR of 1.3), and disorders of lipoprotein metabolism (p = 8.93 x 10^-15^, OR of 1.3), as well as heart failure (p = 1.83 x 10^-22^, OR of 1.95) and other cardiovascular complications (Figure S7D). These data thus suggest circulating γ-synuclein is significantly associated with impairments of cardiometabolic health.

## DISCUSSION

Maintaining metabolic homeostasis requires fine-tuned communication between cell types within a tissue and between distant organs. Disruption of these signals contributes to metabolic and inflammatory disorders. Yet studying intercellular communication *in vivo* has remained challenging due to the lack of tools to precisely identify the tissue of origin of secreted proteins. Here, we present a systematic, tissue-resolved map of ER-resident and secretory pathway proteomes across major metabolic cell types, including adipocytes and hepatocytes at baseline and in response to fasting, LPS treatment, and diet-induced obesity. In addition to defining cell type-specific secreted proteomes, our approach also revealed alterations in proteostatic processes in ER, including protein folding and ER stress responses which become disrupted in energy imbalance.^47^

This work was made possible by several key technological advances. First, we captured organ-specific circulating proteomes using a newly generated, genetically encoded PL mouse model with stable, Cre-dependent expression of an ER-targeted TurboID construct. This bypasses the limitations inherent in viral transduction of PL constructs, including inconsistent gene delivery, off-target effects, and inflammatory responses.^48,49^ Our genetic approach facilitates robust and reproducible labeling of proteins within the ER or destined for secretion. Second, our quantitative proteomic framework, based on multiplexed TMT labeling, improves quantitative accuracy and substantially reduces missing values compared to label-free methods,^14,50^ thereby increasing sensitivity for low-abundance proteins. Finally, our bioinformatics pipeline, including ROC-based enrichment analyses^11,15^ and GO pathway mapping, ensured robust identification of bona fide secreted proteins and biologically relevant pathways.

Our data demonstrate that metabolic stress alters ER protein homeostasis in a dynamic and tissue-specific manner. In adipocytes, our findings revealed distinct secretory pathway proteome remodeling in response to metabolic and inflammatory stress during negative energy balance induced by fasting and LPS. Fasting suppressed anabolic lipid metabolism and ECM remodeling, while enhancing lipid mobilization pathways, suggesting adaptive responses to energy deprivation. The link between systemic ECM changes and caloric restriction has been previously demonstrated in a study of the effects of fasting in human subjects and is thought to contribute to the adaptation to reduced caloric intake.^51^

Fasting also reduced ER-resident and secreted immune proteins, reflecting an energy-conserving adaptation under nutrient deprivation^52^ at the cost of immune function. In contrast, LPS induced impaired lipid processing and activation of inflammatory and ER stress pathways, suggesting possible contributions of acute inflammation to metabolic dysfunction. These changes are consistent with TLR4-mediated immune activation in adipocytes^53,54^ and reprogramming of the secretory machinery toward an immunometabolic phenotype to support its activation.

Hepatic responses were equally distinct; fasting suppressed complement production and promoted metabolic adaptations including an increase in proteins crucial for gluconeogenesis, induction of PPARα target genes like CYP4A14 that drive fatty acid ω oxidation,^55^ and LPIN1, a regulator of triglyceride remodeling and lipolysis.^56^ In contrast, LPS treatment amplified NF-κB/STAT3 inflammatory signaling, elevated the hepatic acute-phase response,^57–59^ and induced proteins within the SREBP2-dependent cholesterol biosynthesis axis (SREBF2, HMGCR, SQLE, FDFT1, CYP51A1). This shift supports membrane biogenesis, lipoprotein remodeling, and receptor trafficking during inflammation, consistent with prior reports.^60^ Collectively, these findings highlight two divergent adaptations: fasting activates lipid catabolism, while LPS initiates a distinct inflammatory metabolic shift characterized by cholesterol biosynthesis and acute phase responses.

The tissue-specific secretome atlas reported here provides a comprehensive resource for functional validation and hypothesis generation. Integration with the UKBB revealed strong links among several circulating proteins and cardiometabolic diseases, such as hypertension, hyperlipidemia, coronary heart disease, and T2D. This included G6PE which was elevated in both inguinal adipocytes in early obesity and adipo-plasma during fasting and early and late-stage obesity. This protein has a role in cell growth, redox balance and ER stress regulation, and its deficiency correlates with fasting hypoglycemia, enhanced insulin sensitivity,^61^ and increased renal oxidative stress.^62,63^ G6PE has not been previously annotated as a circulating factor, however its secretion by adipocytes opens new avenues for understanding its influence on systemic metabolic regulation. G6PE was not included for detection in the ∼3,000 protein Olink panel used in the UKBB cohort, thus the link between circulating G6PE and human disease is currently not known.

We also found that GPR50, an orphan receptor, showed a striking 30-fold induction in adipocytes, particularly visceral fat cells, during diet-induced obesity. While prior studies have linked this X chromosome gene to triglyceride and HDL regulation in obese humans,^43^ the precise cellular function of GPR50 has remained elusive. GPR50 knockout mice exhibit a complex metabolic phenotype and our findings suggest a previously unappreciated role for GPR50 in adipocytes during obesity warranting further investigation.^64^

Diet-induced obesity also led to a dynamic reprograming of the adipocyte proteome. These changes included time-dependent suppression of tissue-resident or locally secreted proteins linked with ECM organization and altered levels of proteins linked with immune system processes, protein folding, and ER stress. We also observed a time- and fat depot-dependent increase of three proteins previously described as neurotrophic factors when obesity develops: CDNF, MANF, and NENF. CDNF is known for its role to negatively regulate pro-inflammatory cytokine secretion and ER stress / UPR activation.^65^ We found that CDNF was induced in inguinal adipocytes but reduced in visceral adipocytes after 15 weeks of HFD. Consistent with this depot-dependent CDNF expression pattern, we identified additional changes in proteins involved in ER stress, including negative regulators of the UPR and integrated stress response in iWAT compared to eWAT cells. MANF has previously been demonstrated to promote iWAT browning and reduce WAT inflammation and hepatic steatosis in obese mice.^66^ Lastly, NENF is a neurotrophic factor known to regulate energy balance via the hypothalamic-peripheral axis^67^ and to modulate sympathetic neuron activity. This is consistent with our data showing that its expression is altered both during inflammation-induced anorexia^68^ and obesity. Collectively, our data suggest that iWAT may modulate its own stores of these neurotrophic factors depending on nutritional intake and energy status.

Finally, our study establishes SYUG as an adipocyte-derived circulating factor, whose plasma levels are acutely and chronically modulated in response to nutritional status. SYUG is elevated in chronic obesity and diminished during fasting or LPS exposure. Although historically studied in neuronal contexts, a role for SYUG has been recently implicated in adipocyte lipolysis via interactions with ATGL and SNARE complex formation.^46,47^ Indeed, independent studies have shown SYUG is a target of peroxisome proliferator-activated receptor gamma 2 (PPARγ2)^69^, the master regulator of adipogenesis, suggesting SYUG is an actively transcribed adipocyte-enriched gene. Notably, whole-body SYUG knock-out mice were protected from DIO and its diabetic complications via increased energy expenditure and improved insulin signaling as well as lower deposition of TGs in liver. Enrichment of SYUG mRNA levels in human WAT in obesity, together with strong associations between basal circulating SYUG levels and cardiometabolic diseases in the UKBB cohort, further underscore its potential role as a significant fat-derived endocrine mediator.

In summary, this study provides a comprehensive resource for elucidating proteomic signatures across physiological states and reveals organ-specific responses to metabolic perturbations, providing critical insights and hypotheses to test into the mechanisms governing metabolic homeostasis. Together, these findings represent an important advance toward identifying novel biomarkers and therapeutic targets for metabolic disorders.

### Limitations of the study

Collectively, our research introduces a powerful genetic and proteomic framework for dissecting the complexities of organ-specific secretomes *in vivo*. However, there are several limitations worth noting. First, the breadth of MS coverage is influenced by multiple factors, including TurboID expression levels, which can vary depending on Cre-driver lines, and the inherent heterogeneity of cell populations within organs. Second, detecting extremely low-abundance proteins or polypeptides derived from non-canonical open reading frames remains challenging and will require further improvements in mass-spectrometry instruments and analysis pipelines. Furthermore, improvements in multiplexing technologies, including the use of recently developed 32-plex TMT reagents, may further enhance experimental throughput and statistical power for future comparative proteomic analyses. Critically, while our proteomic data effectively pinpoint candidate proteins and pathways associated with various metabolic states, rigorous functional studies are indispensable to establish causal relationships between specific secretory changes and observed metabolic phenotypes.

While mechanistic studies are imperative to elucidate the roles of SYUG, GPR50, and G6PE in metabolic homeostasis, this resource will serve as a catalyst for further research. Additionally, expanding the proteomic atlases to encompass additional metabolically active tissues (e.g., pancreas, brain, skeletal muscle) and diverse cell types will generate a more comprehensive understanding of metabolic regulation in pathophysiology. The versatile PL platform developed here can be readily adapted to investigate cell-to-cell communication in other disease contexts or physiological states. Future research will also prioritize dissecting the intricate crosstalk between different cell types and identifying sites and mechanisms of action for newly characterized secreted factors.

## AUTHOR CONTRIBUTIONS

Conceptualization: KHL, EVV, PC, KP; Construct generation: KHL, JMF; Animal experiments: KHL, KP, CRW, BM, SMA, NGB, KM; Mass spectrometry: EVV, KHL, CRW, KP, SMA; Western blot experiments: KP, BM; Bioinformatic analyses: HS, CRW, NR, EVV; Resources: EVV, PC, KHL, JMF; Funding acquisition: EVV, PC, KHL, JMF, KP, CRW; Supervision: EVV, PC, KHL; Writing: KP, CRW, KHL, BM, HS, PC, EVV; Revising: EVV, PC, KHL, JMF.

## ACKNOWLEDGEMENTS

K.P. was funded by the Novo Nordisk Foundation Postdoctoral Fellowship (NNF19OC0055021) and the Elizabeth Hall Janeway Award from the Kellen Women’s Entrepreneurship Fund. C.R.W. was supported by the Marlene Hess Center for Research on Women’s Health and Biomedicine. H.S. and C.R.W. were also supported by Weill Cancer Hub East. B.M. and K.L. were supported by funding from the NIH (P30DA018343 and R00DK129712). P.C. was supported by NIH RC2 DK129961 and the Leducq Foundation. E.V.V. was supported by Robertson Foundation, The Achelis and Bodman Foundation, and Searle Scholar Program. S.M.A. and E.V.V. were also supported by Rockefeller University start-up funds and Irma T. Hirschl/Monique Weill-Caulier Award. Some analyses were performed on computational resources from the Rockefeller University High Performance Computing Resource Center, RRID: SCR_025889.

## RESOURCE AVAILABILITY

### Contact for reagent and resource sharing

Requests for additional information, resources, or reagents should be directed to the lead contact, Ekaterina V. Vinogradova, who will oversee their fulfillment. Questions and requests for mouse models generated in this study should be directed to Ken H. Loh and Paul Cohen and will be fulfilled upon completion of a material transfer agreement.

### Data and code availability

Raw proteomic data and the code used for data processing and visualization will be deposited to PRIDE and Github. Any additional information required to reanalyze the data reported in this paper is available from the lead contact upon request.

## Supplementary information

### (A) Supplementary figures

**Figure S1.**
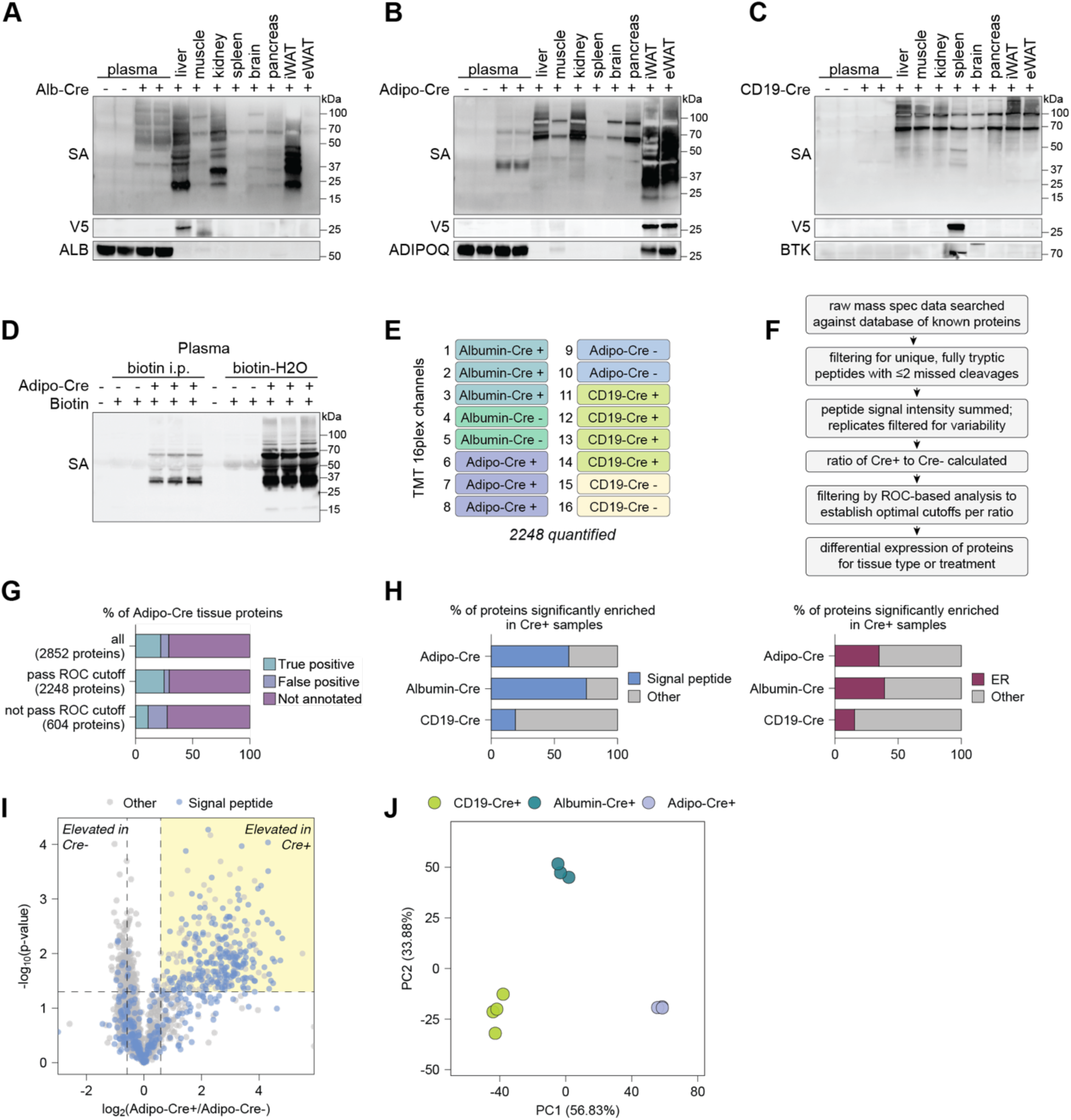
Characterization of a genetically encoded TurboID-KDEL in three tissue types, related to Figure 1. (**A-C**) Protein biotinylation and V5 expression in various tissue types and plasma assessed by Western blot analysis in (A) Albumin-TurboID^KDEL^, (B) Adiponectin-TurboID^KDEL^, and (C) CD19-TurboID^KDEL^ mice. (D) Comparison of protein biotinylation efficiency in plasma from Adipo-TurboID^KDEL^ mice treated with biotin (1 mg/mL) either intraperitoneally (i.p.) or in drinking water over the course of 7 days. (E) Representative TMT channel assignments for a 16-plex LC-MS/MS/MS experiment comparing three tissue types under basal conditions. The number of proteins passing the ROC cutoff is also shown. (F) General filters used for analysis of tissue mass spectrometry data. (G) Representative bar plot showing annotation of proteins passing and failing ROC-based filtering in Adipo-Cre samples. See supplementary methods for more details. (H) Bar plot showing percentage of significantly enriched proteins (p-value < 0.05, FC > 1.5; corresponding to the highlighted region in I) annotated as secreted by SignalP and Outcyte (left) or ER by Gene Ontology annotation (right). (I) Representative volcano plot showing log_2_ fold changes of protein expression between Adipo-Cre+ and Adipo-Cre- samples. Dashed lines represent cutoffs of p-value < 0.05 and fold-change (FC) > 1.5. Signal peptides are labeled based on OutCyte predictions. (J) Principal component analysis of proteomic data from Cre+ samples. Protein signal intensity values were log_2_ transformed before analysis.

**Figure S2.**
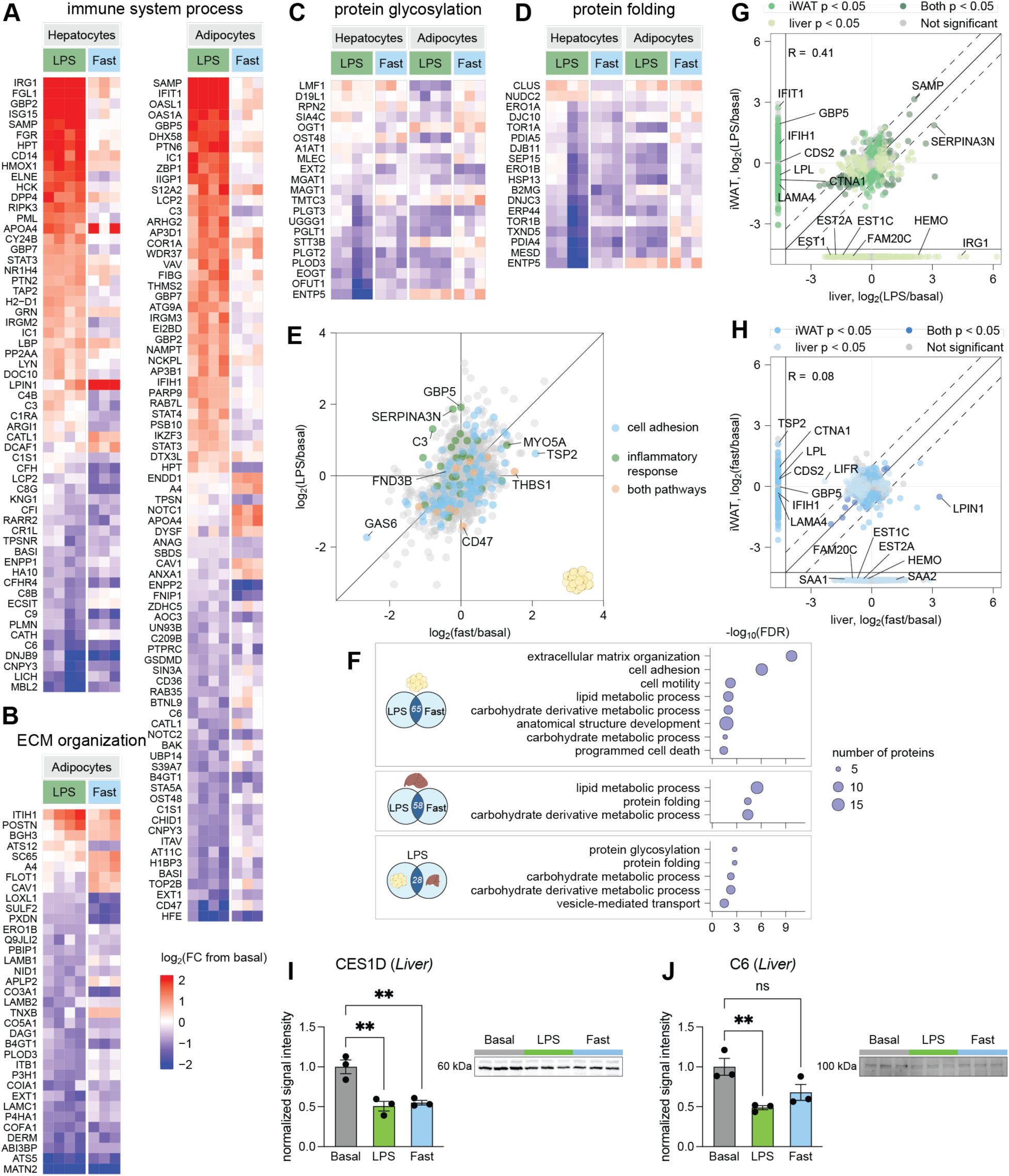
Comparison of proteomic responses to negative energy balance in hepatocytes and adipocytes, related to Figure 2. (**A-D**) Heat maps showing differentially expressed proteins contributing to (A) immune system process, (B) extracellular matrix (ECM) organization, (C) protein glycosylation, and (D) protein folding GO-term enrichment in adipocytes or hepatocytes. Data is presented as log_2_(fold-change) compared to basal condition. (E) Scatterplot showing correlation of log_2_(fold-change) values for protein expression changes in adipocytes from LPS-treated or fasted Adipo-TurboID^KDEL^ mice relative to basal conditions. Proteins belonging to selected pathways are colored. (F) Gene ontology (GO) analysis of pathways enriched at the intersection of proteins significantly downregulated in multiple conditions, including 66 proteins downregulated in adipocytes following LPS and fasting, 58 downregulated in hepatocytes in LPS and fasting, and 28 downregulated in both adipocytes and hepatocytes following LPS. (**G-H**) Scatterplots of log_2_(fold-change) values for protein expression changes in liver (x) or iWAT (y) following LPS treatment (G) or fasting (H) compared to basal condition. (**I-J**) Validation by Western blot of CES1D (I) and C6 (J) downregulation in response to LPS and fasting in liver. Western blot band intensity was normalized to the total intensity of the corresponding lane in a stain-free gel image. Comparison performed by one-way ANOVA with Dunnett’s multiple comparisons test (ns, p > 0.05; **, p < 0.01).

**Figure S3.**
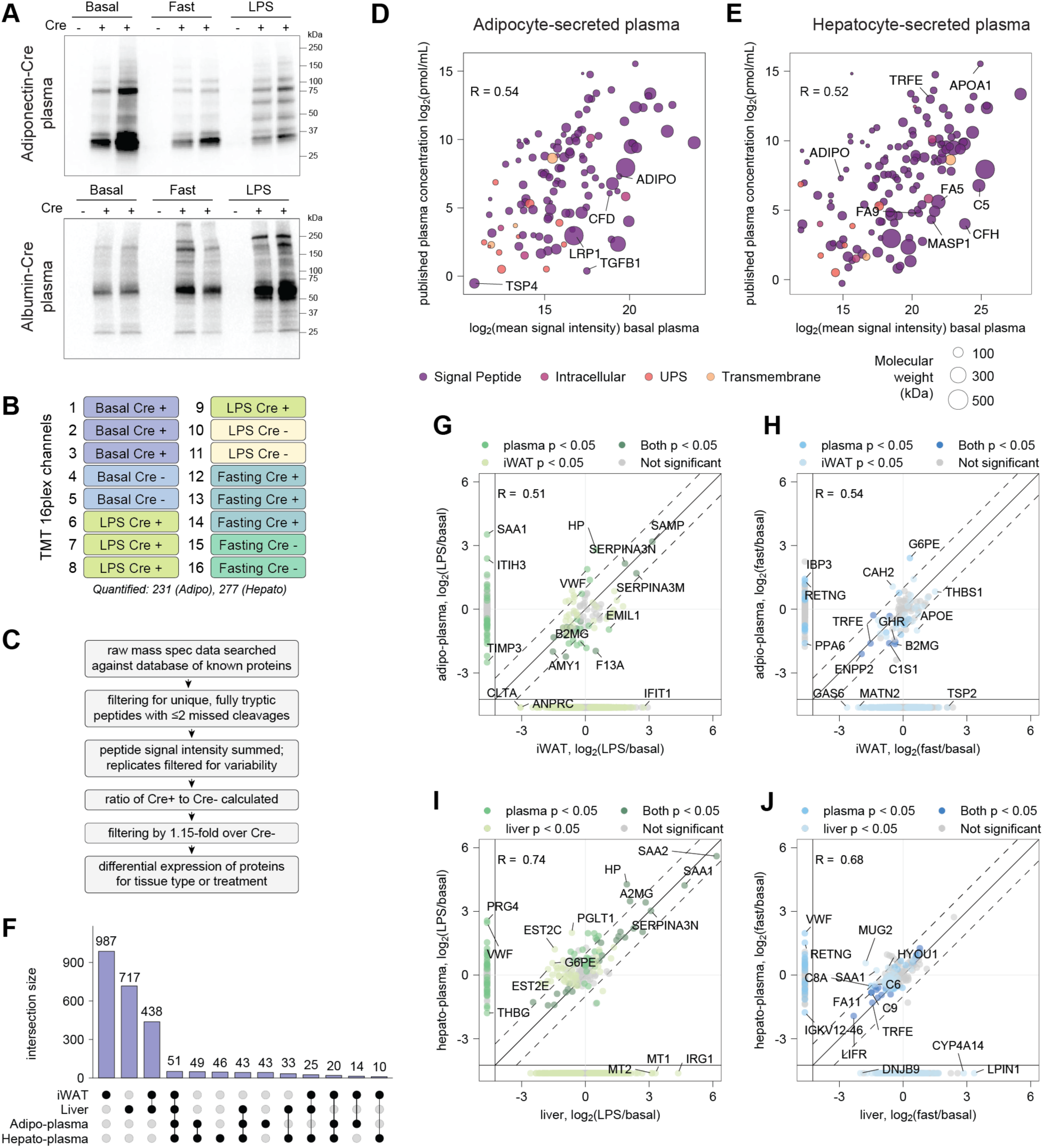
Hepato-plasma and adipo-plasma proteome responses to fasting or inflammation-induced anorexia, related to Figure 3. (A) Western blot showing TurboID-catalyzed biotinylation of proteins identified in plasma from biotin-treated Adipo- and Albumin-TurboID^KDEL^ mice under basal and negative energy balance (fasting, LPS) conditions. (B) Representative TMT channel assignment for a 16-plex LC-MS/MS/MS experiment comparing plasma proteomes from three conditions (basal, LPS, and fasting) in mice with and without a cell-type-specific Cre driver. The number of proteins passing the enrichment cutoff is also shown. (C) Filters used for analysis of plasma mass spectrometry data. (**D-E**) Scatterplots of log_2_-transformed mean signal intensity values (x) for proteins quantified in basal adipo-plasma (D) and hepato-plasma (E) versus mean published absolute plasma concentrations (y). Size of points represents molecular weight in kilodaltons. Colors show SignalP and Outcyte secretion predictions. (**F**) Upset plot of overlap in protein quantifications by TurboID-TMT between adipocyte and hepatocyte tissues and plasma. Intersections with fewer than 10 proteins are not shown. (**G-J**) Scatterplots of log_2_(fold-change) values for expression changes in adipocyte (F-G) or hepatocyte (H-I) secreted proteins following LPS treatment (left) or fasting (right) compared to basal condition in plasma (x) and tissue (y) samples. Proteins quantified in one sample (tissue or plasma) are shown on margins. R represents Pearson’s correlation coefficient.

**Figure S4.**
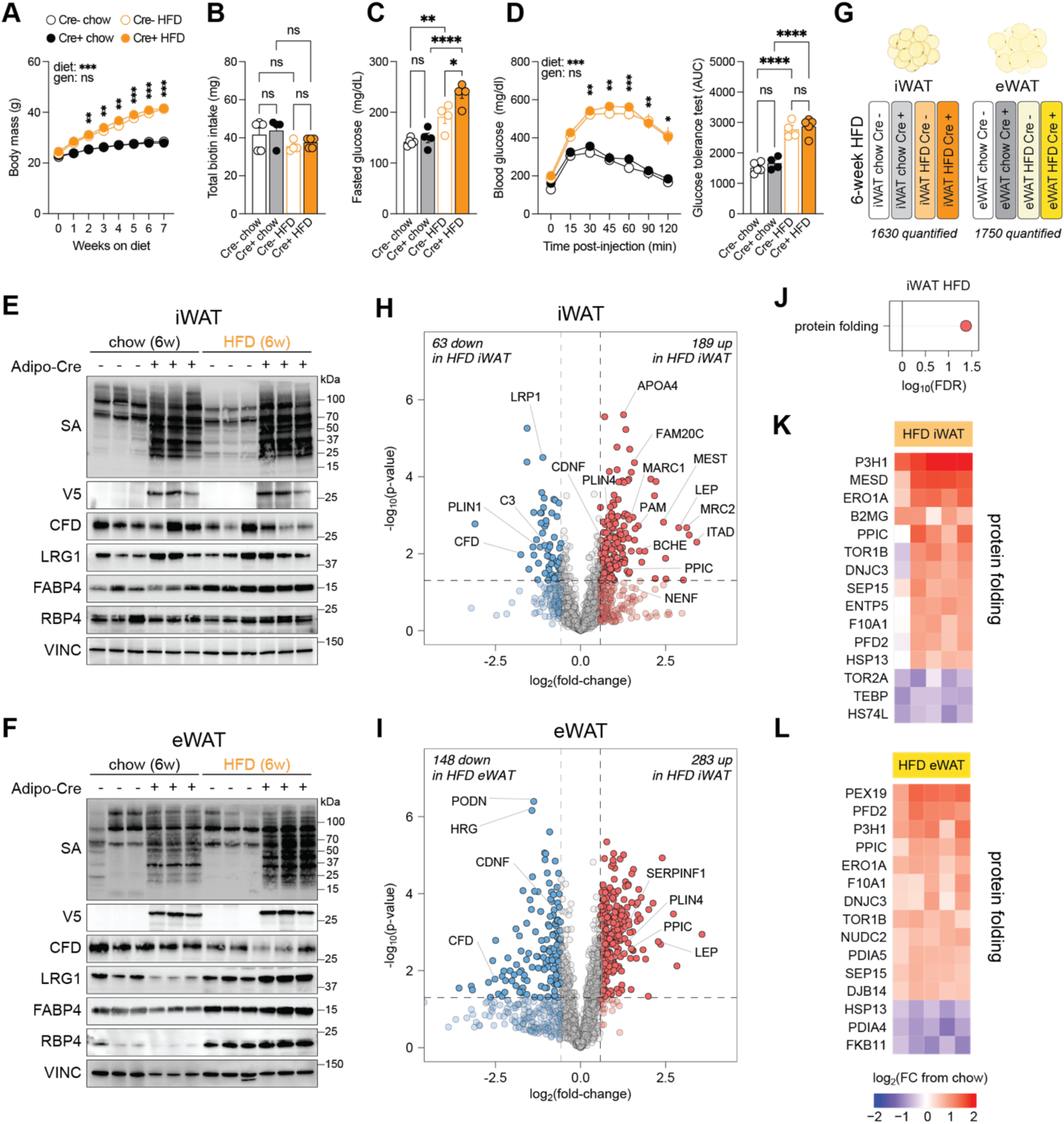
Proteomic responses of inguinal and epidydimal adipocytes in Adipo-TurboID^KDEL^ mice to early obesity, related to Figure 4. **(A)** Body weights of Adipo-TurboID^KDEL^ mice fed HFD or chow for 6 weeks. Biotin (1.5 mg/mL) was administrated in drinking water for 7 days on the last week of feeding. Conditions were compared to basal by two-way ANOVA with Tukey’s multiple comparisons test (ns, p > 0.05, not shown; *, p < 0.05; **, p < 0.01). Error bars show mean ± SEM; n = 4-6 mice per condition. (**B**) Total biotin intake in mice after 7 days. Conditions were compared to basal ordinary one-way ANOVA with Šidák multiple comparisons test (ns, p > 0.05). Error bars show mean ± SEM; n = 4-7 mice per condition. (**C**) Fasted glucose in mice fed chow or HFD for 6 weeks. Conditions were compared to basal ordinary one-way ANOVA with Šidák multiple comparisons test (ns, p > 0.05; *, p < 0.05; **, p < 0.01; ****, p < 0.0001). Error bars show mean ± SEM; n = 4-7 mice per condition. (**D**) Glucose tolerance in mice fed chow or HFD for 6 weeks and area under curve (AUC). Conditions were compared to basal by two-way ANOVA with Tukey’s multiple comparisons test (ns, p > 0.05, not shown; *, p < 0.05; **, p < 0.01; ***, p < 0.001). Error bars show mean ± SEM; n = 4-7 mice per condition. AUC was compared using ordinary one-way ANOVA with Šidák multiple comparisons test (ns, p > 0.05; ****, p < 0.0001). Error bars show mean ± SEM; n = 4-7 mice per condition. (**E-F**) Western blot validation of protein biotinylation using streptavidin-HRP, tissue-specific V5 expression, and levels of known adipokines including CFD, LRG1, FABP4, and RBP4 in bulk iWAT (E) and eWAT (F) from Adipo-TurboID^KDEL^ mice after 6 weeks of HFD feeding. Vinculin (VINC) was used as a loading control. (**G**) Representative TMT channel assignment for a 16-plex LC-MS/MS/MS experiment comparing proteomic changes induced by 6-week HFD vs. chow in iWAT or eWAT adipocytes from Adipo-TurboID^KDEL^ Cre+ and Cre- mice (technical replicates not shown). The number of proteins passing the ROC cutoff in each experiment is also shown. **(H-I)** Volcano plots showing log_2_ fold changes in secretory pathway protein expression in iWAT (H) and eWAT (I) samples following HFD feeding for 6 weeks relative to basal. Dashed lines represent cutoffs of p-value < 0.05 and fold-change > 1.5. (**J**) Biological process over-represented in gene set enrichment affected by HFD feeding in iWAT-derived adipocytes at 6 weeks of HFD. (**K-L**) Heat maps showing differentially expressed proteins contributing to protein folding GO-term enrichment in iWAT (K) and eWAT (L) samples following HFD feeding for 6 weeks. Data is presented as log_2_(fold-change) compared to the chow condition.

**Figure S5.**
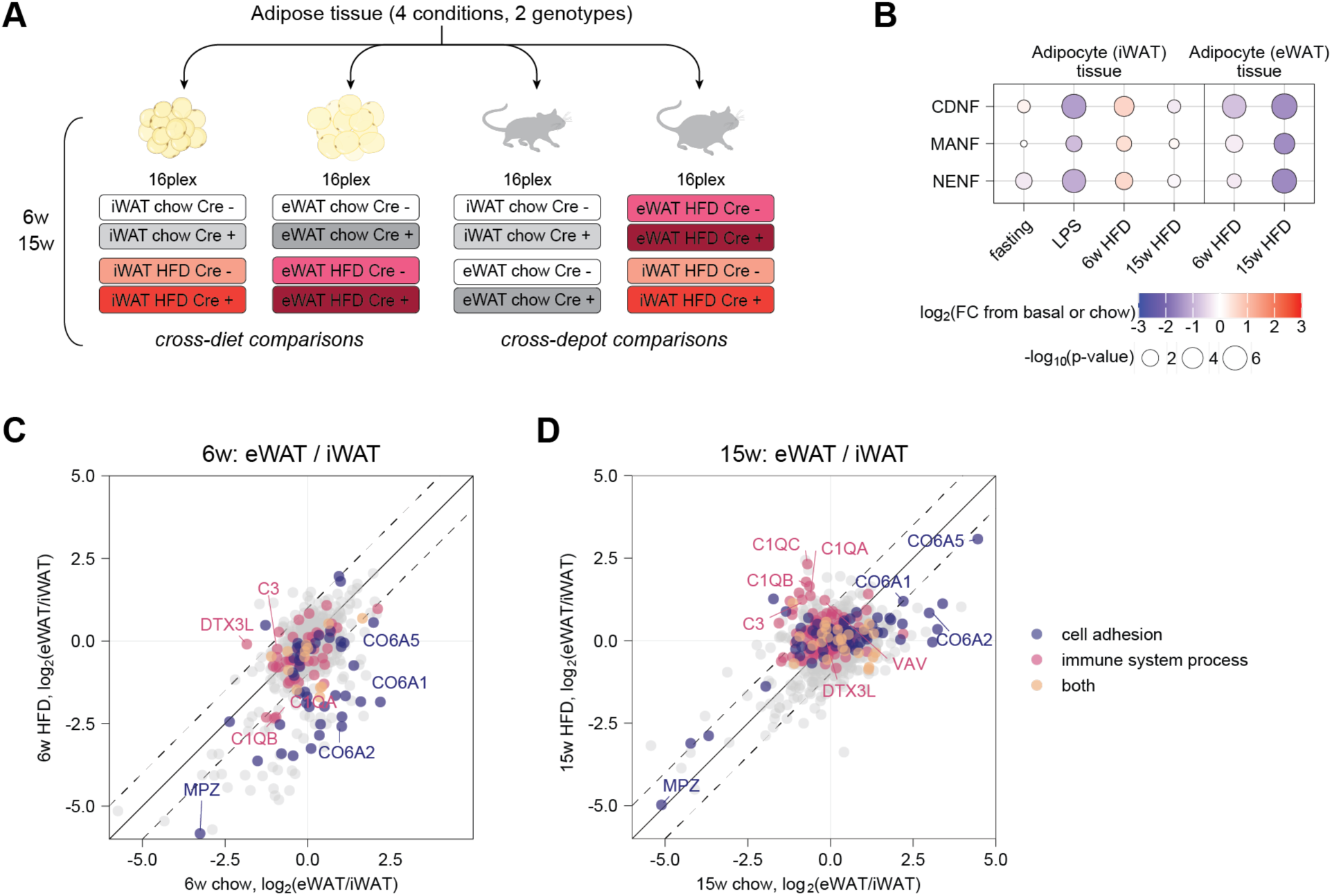
Depot-specific protein remodeling in inguinal and epididymal adipocytes in early and advanced obesity, related to Figure 5. (A) Schematic showing TurboID-TMT MS experiments for cross-diet and cross-depot comparisons (technical replicates not shown). (B) Normalized mean signal intensity values for CDNF, MANF, and NENF proteins across high-fat diet experiments and depot. (**C-D**) Correlational analysis of log_2_ fold changes in protein expression following chow and HFD feeding for 6 weeks (C) and 15 weeks (D) in inguinal compared to epididymal adipocytes. Proteins belonging to selected pathways are colored.

**Figure S6.**
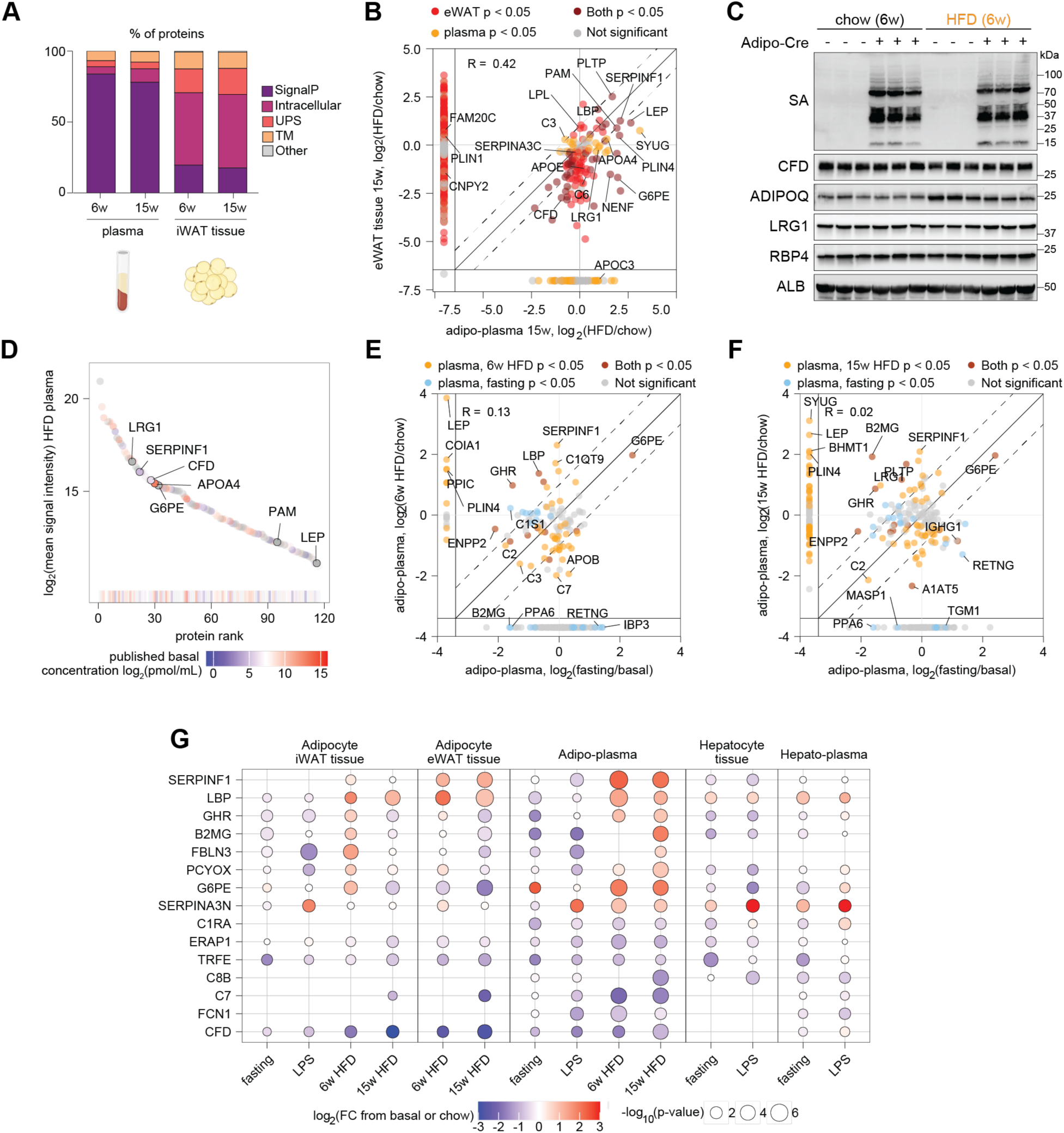
Characterization of adipo-plasma proteome during obesity, related to Figure 6. (A) Bar plot of proteins quantified in Adipo-TurboID^KDEL^ iWAT and plasma proteomes at 6 and 15 weeks of DIO, with protein class labeled based on SignalP and OutCyte predictions. (B) Scatterplot of log_2_ fold changes in protein expression in eWAT vs. plasma proteomes of Adipo-TurboID^KDEL^ mice following HFD feeding for 15 weeks compared to chow. Proteins quantified for only one of the conditions are shown on margins. (C) Western blot validation of protein biotinylation using streptavidin-HRP and levels of known adipokines including CFD, ADIPOQ, LRG1, and RBP4 in bulk plasma samples from Adipo-TurboID^KDEL^ mice after 6 weeks of HFD feeding. Albumin (ALB) was used as a loading control. (D) Rank plot of mean signal intensity values of proteins quantified by TurboID-TMT in Adipo-TurboID^KDEL^ plasma proteome following HFD feeding for 6 weeks colored by known basal plasma concentrations. (**E-F**) Scatterplots of log_2_ fold changes in protein expression in plasma proteomes of Adipo-TurboID^KDEL^ mice following 48-hour fasting and HFD feeding for 6 weeks (E) or 15 weeks (F) compared to basal or chow. Proteins quantified for only one of the conditions are shown on margins. (**G**) Dot plot of log_2_ fold changes and p-values of selected proteins detected in adipocytes, hepatocytes, and plasma across negative and positive energy balance conditions. Data are shown relative to own control group (e.g. fasting vs. basal, LPS vs. basal, HFD vs. chow).

**Figure S7.**
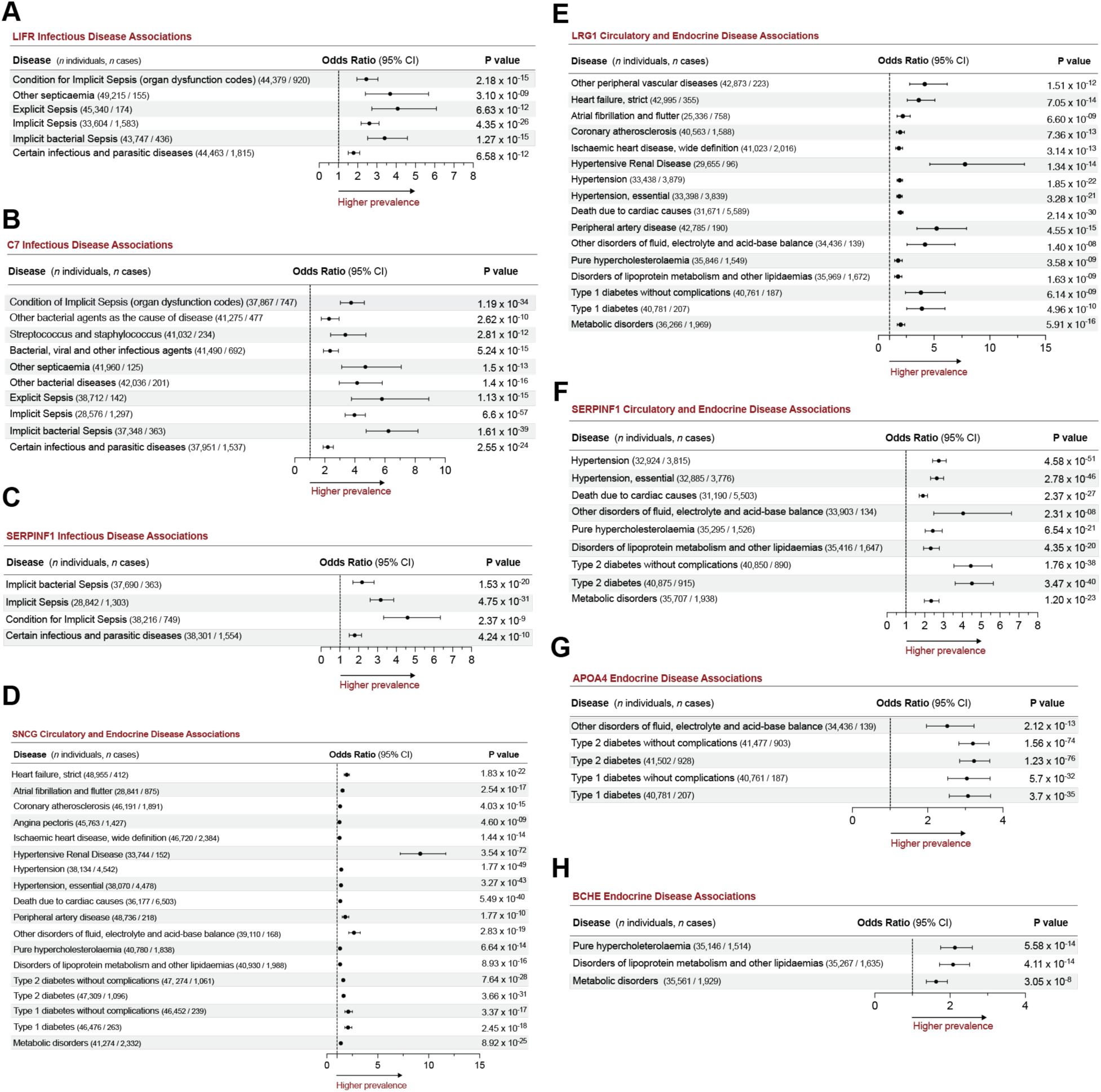
Odd ratios for circulatory, endocrine, and infectious disease and clinical outcomes linked with individual circulating proteins derived from adipocytes or hepatocytes in various energy balance states, related to Figure 7. (**A-H**) Odds ratios for infectious diseases, endocrine, and/or circulatory prevalence-based outcomes linked with basal circulating levels of leukemia inhibitory factor receptor (LIFR), complement component 7 (C7), serpin family F member 1 (SERPINF1) protein, gamma synuclein (SNCG), leucine-rich glycoprotein 1 (LRG1), apolipoprotein A-IV (APOA4), and cholinesterase (BCHE) in Health and Disease Atlas based on UKBB cohort of 53,026 individuals.

### (B) Supplementary Data Set Legends

**Table S1: Experimental metadata and protein filtering metrics.**

Details of mouse biotin labeling experiments, mass spectrometry settings, and protein quantification statistics. An ROC filter was applied for tissue experiments, while a 15% enrichment filter was applied for plasma experiments. B-cell, hepatocyte, and adipocyte enriched proteins were determined by retrieving genes from the human protein atlas in categories group enriched, cell type enriched, and cell type enhanced.

**Table S2: Mean protein signal intensity across all experiments.**

Signal intensity data is mean of technical replicates. Median normalization was performed for cre+ channels for each condition. Cre- channels are unnormalized values from IP2 search. Published plasma concentrations are calculated from Michaud et al 2018.

**Table S3: Differential expression of proteins in the mouse secretome.**

TurboID-TMT data showing protein expression changes in the mouse secretome. For every protein the log_2_ fold-change of the ratio between condtions was calculated. p-values were calculated with T-test for the means of two independent samples. Infinite log_2_(fold-change) values were replaced with the greatest non-infinite value in each direction (10.197 and -8.627)

**Table S4: Proteome-phenome atlas assocations.**

Protein disease associations for proteins quantified in the mouse secretome. Associations were retrieved from the proteome phenome atlas. Associations are filtered to prevalent associations with Bonferroni-corrected p-value < 0.05, in chapters I, IV, and IX.

### (C) Experimental model and subject details

#### Mice

Experiments involving mice were approved by The Rockefeller University’s Institutional Animal Care and Use Committees (Protocol numbers 21037, 19087, and 24019-H), and Yale University’s Institutional Animal Care and Use Committee (Protocol number 2022-20445).

Mice were housed 2-5 per cage in a 12-hour light/12-hour dark cycle with *ad libitum* access to water and regular chow except in fasting studies where chow was restricted. We used WT male C57BL/6J mice (Jackson Laboratory 000664), Rosa26-Cas9 knockin (*Gt(ROSA)26Sor^tm1.1(CAG-cas9*,-EGFP)Fezh^*/J, Jackson Laboratory 024858) Albumin-Cre (B6.Cg-Speer6-ps1^Tg(Alb-cre)21Mgn^/J, Jackson Laboratory 003574). CD19-Cre (B6.129P2(C)-Cd19tm^1(cre)Cgn^/J, Jackson Laboratory 006785), Adiponectin-Cre (B6.FVB-Tg(Adipoq-cre)1Evdr/J, Jackson Laboratory 028020), and LSL-TurboID-KDEL, which was generated in the transgenic core facility at Rockefeller University. All mouse lines are on a wild type (C57BL/6J) background. Littermates of the same sex (all male) were randomly assigned to either experimental or control groups.

#### Construct generation and genotyping

LSL-V5-TurboID-KDEL-IRES-eGFP was cloned by Gibson Assembly into the CTV plasmid (Addgene #15912) at the Asc1 restriction site for recombination into the Rosa26 locus. Transgenic knock-in chimeric mice were generated by ES cell injections into mouse blastocysts, at Rockefeller’s Transgenic and Reproductive Technology Center. Mice were backcrossed eight generations in the C57BL/6J background before additional crosses. Mice were then bred to homozygosity without any obvious developmental defects. Primers for genotyping both homozygous and heterozygous mice were:

TurboID^KDEL^ Fwd 5’ CCC GAG CCT ATC CCG CTG CTG AAC 3’

TurboID^KDEL^ Rev 5’ CCC CAA TGA TCA GCT TCA CGG GTC 3’

Rosa26 Fwd 5’ AAA GTC GCT CTG AGT TGT TAT 3’

Rosa26 Rev 5’ GGA GCG GGA GAA ATG GAT ATG 3’

#### Biotin water

Biotin-containing water for basal profiling and negative energy balance experiments was prepared by dissolving solid biotin in sterile drinking water at a concentration of 0.25 mg/mL. To solubilize a higher concentration of biotin for HFD experiments (1.5 mg/mL), biotin was first dissolved in a small volume of Trizma base (3 g biotin, 20 mL, 1 M Trizma base) and then diluted in sterile drinking water to 2 L final volume with pH returned to pH 7.5 by addition of hydrochloric acid. Mice were provided *ad libitum* access to biotin-containing water during treatment.

#### LPS treatment

Heterozygous (TurboID^KDEL^/C57BL/6J) mice, with and without expression of Adiponectin- or Albumin-Cre, were used for LPS and fasting experiments. For LPS-treated animals, a frozen aliquot of LPS (Sigma Aldrich L2880, 1 mg/mL) was diluted in sterile phosphate-buffered saline (PBS) to a final concentration of 0.1 mg/mL. This solution was injected into each animal i.p. at a concentration of 1 mg/kg while saline alone was provided as a control.

#### Food restriction & chow consumption measurements

For fasted animals, food was removed from the hopper for 48 hours, while other groups of animals had *ad libitum* access to chow. Food intake was measured daily.

#### Diet-induced obesity

Homozygous (TurboID^KDEL^/TurboID^KDEL^) mice crossed with Adiponectin-Cre donors^1^ were used for diet-induced obesity studies. All mice were group-housed (3-4 per cage), maintained at RT in 12-hour light:dark cycle (0700-1900) and allowed *ad libitum* access to water and food (chow or 60 kcal% high-fat diet, see STAR Methods Table). For early-stage obesity, male mice of both genotypes (Adipo-TurboID^KDEL^ Cre+ and Cre-) were weighed, randomly allocated to diet groups (minimum n=4 per condition) and terminated after 6 weeks of HFD feeding. An independent age-matched cohort fed HFD/chow for 6 weeks was used for glucose tolerance tests and body composition scans. For late-stage obesity, male mice were fed HFD/chow for 15 weeks starting at 7 weeks of age. Biotin was administered in drinking water for 7 days prior to plasma and tissue harvests. Biotin intake was recorded daily, and body weights were recorded weekly.

#### Metabolic phenotyping in diet-induced obese mice

For fasting glucose measurements and glucose tolerance tests (GTT) mice were housed individually and fasted for 6 hours in a noise-free procedure room. Circulating levels of glucose were determined from tail-derived blood using a standard glucometer. Following fasting glucose at time 0, mice were injected i.p. with bolus glucose at 1.5 g/kg in 30-second intervals and glucose measurements were taken at 15, 30, 60, 90, and 120 min post-injection. Two measurements with two separate glucometers were taken per mouse and per timepoint to ensure measurement accuracy (means are reported). Body composition (fat mass, lean mass) was assessed using rodent echoMRI.

#### Blood glucose measurements

A small incision was made on the mouse’s tail to allow collection of blood, and blood glucose levels were measured using a Contour NextEZ glucometer.

#### Sample collection

Plasma sample collection: Mice were anesthetized with an overdose of isoflurane and blood was collected via cardiac puncture, using EDTA coated microcentrifuge tubes (Sarstedt), and placed on ice. Tubes were centrifuged at 1,000 *g* for 20-30 min at 4°C, followed by collection of the plasma fraction and storage at - 80°C until further analysis.

Tissue sample collection: Following blood collection, mice were perfused with PBS to minimize any non-specific residual biotinylation signal, organs were then dissected out, rinsed with ice-cold PBS briefly, snap frozen in liquid nitrogen, and stored at -80°C before use.

Fixed tissues for validation in sections: Mice were anesthetized with isoflurane and an intracardiac perfusion and fixation was performed with 1X PBS followed by cold 4% PFA. All harvested samples were post-fixed in 4% PFA at 4°C for 24 h. Fixed tissue samples were washed with PBS for three times before subsequent steps.

### (D) Experimental methods details

#### Plasma processing

For mass spectrometry, 200-400 µL of plasma was diluted to 15 mL using 1x PBS and centrifuged at 4,000 rpm for 1 h at 4°C using a 3 kDa filter (Millipore, UFC900324) twice to remove biotin, until the solution was concentrated to 0.5 mL. The final 0.5 mL solution was transferred to a new 1.5 mL tube and 0.8 mL of RIPA buffer containing protease inhibitors was added before further analysis or subsequent enrichment steps.

#### Tissue lysis for liver and spleen

Half of a single lobe of liver (∼400 mg) and entire spleen from each animal were used for analysis. Tissues were minced using scissors and one 3 mm Tungsten carbide metal bead was added to the sample in a 2.0 mL microcentrifuge tube, containing 1 mL of cold RIPA buffer (10 mM Tris, 150 mM NaCl, 1% Triton X-100, 1% sodium deoxycholate, 0.1% SDS, pH 7.5) containing cOmplete protease inhibitor cocktail (Roche, 04693116001) and 1 mM PMSF. Tissues were then lysed using a Qiagen TissueLyser II (20 frequency/sec for 20 min). Samples were incubated in RIPA at 4°C with end-over-end rotation for 20-30 min to boost protein extraction yield, followed by centrifugation at 14,000 rpm for 10 min, and collection of the supernatant. Protein concentration was measured using standard DC or BCA assay and subjected to further analysis or enrichment steps described below.

#### Protein extraction from adipose tissues

Adipose tissues (iWAT, eWAT) were harvested from anaesthetized, PBS-perfused mice and snap frozen in liquid nitrogen, then pulverized to a fine powder (Cellcrusher). Soluble proteins were extracted using NP-40-based RIPA buffer (1% NP-40, 150 mM NaCl, 0.5% deoxycholate, 0.1% SDS, 50 mM Tris-HCl pH 7.5) containing cOmplete protease inhibitor cocktail (Roche, 04693116001), and phosphatase inhibitor (Roche, 4906845001) tablets (1 tablet per 10 mL). Lysates were incubated with end-over-end rotation for 30 min at 4°C, followed by centrifugation at 8,000 rpm for 20 min at 4°C, and lipid-free soluble infranatant fractions were extracted for downstream analyses.

#### Immunoblotting

##### Protocol 1 (Figures 1, S1, 4-7)

Protein concentrations were determined using Pierce BCA assay and normalized to 3 µg/µL (iWAT, eWAT) or 9 µg/µL (plasma) in XT sample buffer, boiled at 90°C for 5 min and resolved at 30 µg/well (tissue extracts) or 90 µg/well (plasma) in 12-well 4-12% Criterion XT Bis-Tris gels in MES electrophoresis buffer. Proteins were transferred onto 0.45 µm (tissues) or 0.22 µm (plasma) ImmobilonP PVDF membranes in Tris-Glycine-MeOH buffer at 30V (4°C, overnight), blocked in biotin-free SuperBlock for 1 h at RT, and incubated with Streptavidin-HRP (SA), or primary antibodies against V5 tag, CFD, LRG1, RBP4, FABP4, ADIPOQ, ALB, and VINC, as detailed in the Key Resources Table. HRP-conjugated secondary antibodies were used for detection with enhanced chemiluminescence in ChemiDoc Imager. Tissue panel blots for V5 were run using multiple organs at the same concentration per well (30 µg total) to allow for side-by-side comparison. Vinculin (tissues), Albumin (plasma) or Coomassie Blue were used as loading controls.

##### Protocol 2 (Figures 2 and 3)

Protein samples were combined with SDS protein loading buffer, boiled for 5 min, and resolved by SDS-PAGE. Proteins were transferred onto a nitrocellulose membranes (0.2 μm, Bio-Rad, 1620112), which were then blocked with Odyssey blocking buffer (LICOR) or 3% w/v BSA in TBST (0.1% Tween-20 in Tris-buffered saline) for 30 min. Membranes were incubated overnight at 4°C with primary antibodies diluted in 3% (w/v) BSA in TBST, followed by four 5-min washes in TBST. Following primary antibody incubation, membranes were probed with IRDye 800CW or IRDye 680RD (LICOR Biosciences) (1:10000 dilution) or HRP-conjugated secondary antibodies at RT for 1 h, then imaged using Bio-Rad ChemiDoc. Blots were quantified using Bio-Rad Image Lab software. HRP-labelled blots were developed using a chemiluminescence substrate (Clarity Western ECL Substrate, Bio-Rad, 1705061). Alternatively, samples were imaged using the same channels on a Bio-Rad ChemiDoc.

For Western blot validation (Figures S2I, S2J, 3G), proteins were resolved on 4-20% Mini-PROTEAN TGX stain-free gel (Bio-Rad, 4568095). Western blot band intensity was normalized to the total intensity of the corresponding lane in a stain-free gel image.

#### Immunofluorescence

Adipose, liver or spleen tissue was collected from anesthetized mice perfused with PBS and 4% paraformaldehyde. Liver and spleen were sunk with 30% sucrose in PBS overnight. Adipose tissue was delipidated as previously described.^2^ Samples were embedded in OCT and cut into 30 µm sections at -20°C. Sample sections were blocked in 3% BSA, 1% donkey serum containing PBST (0.1% Triton X-100, 0.05% Tween-20), followed by incubation in primary antibody dilutions for 2 h at RT. After incubation with primary antibodies, samples were washed in PBST four times and then incubated in secondary antibody dilutions for 4 h at RT in blocking buffer. Samples were then washed in PBST four times and mounted with DAPI Fluoromount (Southern Biotech).

#### Confocal microscopy

Representative sample regions of iWAT were imaged using an inverted Zeiss LSM 980 laser scanning confocal microscope with a 40X objective lens. Liver and spleen were imaged using an inverted Leica TCS SP8 X laser scanning confocal microscope with a 40X objective lens.

#### Biotinylated protein enrichment

To enrich biotinylated proteins, 100 μL of streptavidin (SA) magnetic beads (Pierce, 88816) per sample was washed twice with RIPA buffer (150 mM NaCl, 1% NP-40, 0.5% sodium deoxycholate, 0.1% SDS, 50 mM Tris pH 7.5) with protease and phosphatase inhibitors. The washed slurry was then incubated with cell lysates containing the same amount of protein input (3-5 mg of protein) for all samples of that same tissue type and incubated with end-over-end rotation at 4°C overnight. Using a magnetic rack, the beads were subsequently washed twice with 1 mL of RIPA lysis buffer, once with 1 mL of 1 M KCl, once with 1 mL of 0.1 M Na_2_CO_3_, once with 1 mL of 2 M urea (freshly prepared) in 10 mM Tris-HCl (pH 8.0), and twice with 1 mL RIPA lysis buffer. A small volume of protein-bound beads (5% of total volume) was removed at this point for validation by Western blot and bead-bound proteins were eluted by boiling at 95°C in 2X SDS loading buffer containing 20 mM DTT and 2 mM biotin for 15 min.

#### Elution of peptides from beads

To prepare proteomic samples for mass spectrometry analysis, biotinylated proteins bound to SA beads were washed with 1 mL of 50 mM Tris-HCl (pH 7.5), followed by two washes with 1 mL 2 M urea in 50 mM Tris (pH 7.5) buffer. The buffer was removed, the beads were re-suspended in 80 μL of 2 M urea in 50 mM Tris-HCl containing 1 mM DTT, 0.5 mM CaCl_2_, and 0.8 μg trypsin (Promega, V5111), followed by incubation at 25°C for 1.5 h with shaking at 1,000 rpm. After 1.5 h, samples were placed on a magnet, and the supernatant was transferred to fresh low-binding Eppendorf tubes. Streptavidin beads were then washed twice with 60 μL of 2 M urea in 50 mM Tris (pH 7.5), and the washes were combined with the initial supernatant for subsequent reduction, alkylation, and digestion steps.

#### Reduction, alkylation, trypsin digestion, and desalting of peptides

Samples (200 µL total) were reduced with DTT (final concentration: 8 mM; 65°C, 20 min) and alkylated with iodoacetamide (final concentration: 20 mM; 37°C, 30 min in the dark with shaking at 1,000 rpm). An additional 0.5 μg of trypsin was added to each sample and protein digestion was carried out overnight at 25°C with shaking at 700 rpm. After overnight digestion, the sample was acidified to pH 3 by adding formic acid (FA) to a final 1% (vol/vol) FA. Samples were desalted using C18 StageTips (Pierce, 87784). C18 StageTips were first conditioned with 100 µL of 100% methanol, followed by 100 µL of solvent 1 (50:50 acetonitrile (ACN): H_2_O, 0.1% FA), and twice with 200 µL of buffer A (95% H_2_O, 5% ACN, 0.1% FA). Acidified peptides were loaded onto the conditioned StageTips and washed twice with 200 µL of buffer A (95% H_2_O, 5% ACN, 0.1% FA). Samples were eluted from the StageTips with 2 x 200 µL of buffer B (80% ACN/H_2_O, 0.1% FA), and the solvent was removed until completely dry using SpeedVac vacuum concentrator.

#### TMT labeling for multiplexed quantitative proteomics

Peptides were resuspended in 100 µL of 30% ACN in 200 mM EPPS (pH 8.0). Samples were vortexed, spun down, and sonicated in a water bath for 5 min. After brief centrifugation, 3 µL of the corresponding TMTpro 16-plex reagent (Thermo Fisher, A44521) was added to each tube (20 µg/µL in dry ACN). The samples were vortexed, spun down, and left at room temperature for 1 h. To quench the labeling, 3 µL of 5% hydroxylamine was added to each sample. Samples were vortexed, spun down, and left at room temperature for 15 min. Samples were then acidified with 5 µL of FA, combined into two low binding 1.5 mL Eppendorf tubes, and dried using SpeedVac vacuum concentrator, leaving gel-like salt residues.

#### Desalting of pooled multiplexed samples

Following TMT labeling, samples were resuspended in 500 µL of buffer A (95% H_2_O, 5% ACN, 0.1% FA). An additional 20 µL of 20% FA was added to ensure the sample remained acidic. The Sep-Pak C18 cartridge was conditioned by adding 1 mL 100% ACN three times and equilibrated using 1 mL buffer A (95% H_2_O, 5% ACN, 0.1% FA) three times. Samples were added slowly at a rate of 1 drop/sec, and the flow through was collected and re-loaded a second time. To desalt the samples, 1 mL of buffer A (95% H_2_O, 5% ACN, 0.1% FA) was added to the cartridge three times. Samples were eluted by adding 1 mL of buffer B (80% ACN, 20% H_2_O, 0.1% FA). The eluate was then dried using SpeedVac vacuum concentrator.

#### Pre-fractionation by HPLC

Dried desalted samples were resuspended in 500 µL buffer A (95% H_2_O, 5% ACN, 0.1% FA) and loaded on a high-pressure liquid chromatography (HPLC) for offline pre-fractionation. Samples were separated on a C18 column by reverse phase HPLC, starting in a high pH buffer containing 10 mM ammonium bicarbonate over an increasing acetonitrile gradient (0-100%). Samples were collected into a deep 96-well plate, pooled across elution times (12 fractions for tissue, 4 fractions for plasma), and then dried using SpeedVac vacuum concentrator. Because the adipo-plasma and hepato-plasma proteomes were expected to be lower in abundance than the intracellular ER-associated proteomes, we pooled HPLC-fractionated samples into four final fractions to optimize coverage and enhance detection of low abundance factors.

#### Liquid chromatography tandem mass spectrometry analysis (LC-MS/MS/MS)

Samples were re-suspended in 10 µL of buffer A and briefly sonicated before analysis by liquid chromatography tandem mass spectrometry (7.5 μL injection) using Orbitrap Eclipse mass spectrometer (Thermo Fisher) with UltiMate 3000 Series Rapid Separation LC system. Peptides were eluted onto a 2 μm, 100 Å, 75μm x 25 cm capillary column at 0.25 μL/min flow rate and separated using the following gradient: 5% LC-MS buffer B (ACN, 0.1% FA) in LC-MS buffer A (H_2_O, 0.1% FA) from 0-10 min, 5%-20% buffer B from 20-120 min, 20%-45% buffer B from 120-140 min, 45%-95% buffer B from 140-145 min, 95% buffer B from 145-147 min, 5% buffer B from 147-149 min, 95% buffer B from 149-151 min, 5% buffer B from 151-160 min).

The following scan sequence was used: (1) MS1 master scan (Orbitrap analysis, resolution 120,000, scan range 400-1,600 m/z, RF lens 40%, normalized AGC Target 250%, automatic maximum injection time, profile mode with enabled dynamic exclusion (repeat count 1, duration 60 s); (2) Selection of top 20 ions for MS2 analysis; (3) MS2 analysis, which consisted of quadrupole isolation (isolation window 0.7 m/z) of precursor ion followed by collision-induced dissociation (CID) in the ion trap (fixed collision energy mode, normalized collision energy 35%, maximum CID activation time 10 ms, Activation Q 0.25). Except for diet-induced obese and chow-fed iWAT and eWAT analyses, all runs used real-time search with a 2017 mouse FASTA database and included static modifications for cysteine carbamidomethylation (Δ Mass 57.0215), lysine-, and N-terminal amine modification with TMTpro 16-plex reagents (Δ Mass 304.2071), as well as variable modifications for methionine oxidation (Δ Mass 15.9949). Maximum missed cleavages were set to 2, with maximum variable modifications per peptide at 1.

#### RNA preparation and quantitative PCR

Total RNA was extracted from white fat tissue, spleen or liver using TRIzol (Invitrogen) along with QIAGEN RNeasy mini kits. For quantitative PCR (qPCR) analysis, RNA was reverse transcribed using the QuantiTect Reverse Transcription Kit (Qiagen). cDNA was used in qPCR reactions containing SYBR-green, fluorescent dye (ABI). Relative mRNA expression was determined by normalizing to the geometric mean of *Ppib* (cyclophilin B) and *Tbp* (TATA-box binding protein) levels using the ΔΔC_t_ method. Primer sequences are:

*Cyclo*/*Ppib* Fwd 5’ GGA GAT GGC ACA GGA GGA A 3’

*Cyclo*/*Ppib* Rev 5’ GCC CGT AGT GCT TCA GCT T 3’

*Tbp* Fwd 5’ GGG TAT CTG CTG GCG GTT T 3’

*Tbp* Rev 5’ TGA AAT AGT GAT GCT GGG CAC T 3’

*Pam* Fwd 5’ CTG GGG TCA CAC CTA AAG AGT 3’

*Pam* Rev 5’ ATG AGG GCA TGT TGC ATC CAA 3’

*Ppic* Fwd 5’ TGA CGG ACA AGG TCT TCT TTG A 3’

*Ppic* Rev 5’ CAG AGC CAC GAA GTT TTC CAC 3’

*SerpinF* Fwd 5’ GCC CTG GTG CTA CTC CTC T 3’

*SerpinF* Rev 5’ CGG ATC TCA GGC GGT ACA G 3’

*Cdnf* Fwd 5’ CTT TTG CGC CGG GTT TTG TAT 3’

*Cdnf* Rev 5’ AGG GAG TTG TAG AAT CGG TCT AA 3’

*Sncg* Fwd 5’ AAA GAC CAA GCA GGG AGT AAC G 3’

*Sncg* Rev 5’ GAC CAC GAT GTT TTC AGC CTC 3’

*Nenf* Fwd 5’ GAA GGG AGT GGT GTT CGA TGT 3’

*Nenf* Rev 5’ GTG TCG TGA GTG AGG TCT GC 3’

*Flst1* Fwd 5’ CAC GGC GAG GAG GAA CCT A 3’

*Flst1* Rev 5’ TCT TGC CAT TAC TGC CAC ACA 3’

*Gpr50* Fwd 5’ AGA GCA ACA TGG GAC CTA CAA 3’

*Gpr50* Rev 5’ GCC AGA ATT TCG GAG CTT CTT G 3’

### (E) Quantification and statistical analysis

#### Statistical analyses

Statistical analyses were performed using GraphPad Prism and R package. Two-way ANOVA followed by Tukey’s multiple comparisons test was applied to determine the statistical differences among groups across different timepoints. Non-repeated measure data were compared to basal with ordinary one-way ANOVA and Šidák multiple comparisons test. Unpaired two-tailed Student’s t test was applied to determine the statistical differences for gene expression and protein levels. The results of statistical analyses for each experiment can be found in corresponding figure legends. All values are reported as mean ± SEM. P-values below 0.05 were considered significant throughout the study.

#### Data processing for proteomic experiments

Raw data files were processed using RAW converter (available at github.com/proteomicsyates/RawConverter) and the resulting MS1, MS2, and MS3 files were uploaded to the Integrated Proteomics Pipeline (IP2). Peptide identification was performed against a reverse concatenated, non-redundant version of the Mouse Uniprot database (release-2017_07) using the ProLuCID algorithm. Static modifications for TMT labeling were set for N-termini and lysine residues (+304.2071 Da for TMT 16-plex). Search results were filtered with DTASelect (v2.0) to achieve a peptide-level false discovery rate below 1%. MS3 reporter ions were used for quantification with a mass tolerance of 20 ppm via the IP2 platform.

#### Filtering and normalization of proteomic data

Experiments were processed individually with the following filters: removal of non-unique peptides, removal of half-tryptic peptides, removal of peptides with more than two internal missed cleavages, removal of peptides with low (< 5,000) average of reporter ion intensities for all pairs of channels, and peptides with high variation between all pairs of channels (coefficient of variation > 0.5). Additionally, keratin proteins and proteins whose human orthologs were quantified in over 200 experiments in the CRAPome^3^ database were labelled as contaminants and excluded from analysis.

Median normalization was performed on signal intensities for Cre+ channels within each experiment. True positive proteins (defined in Receiver operating characteristic analysis) with 2+ peptides were used to calculate the median per channel and normalization factors were calculated by dividing the median of all Cre+ channel medians by each individual channel’s median. Each Cre+ channel was then multiplied by its corresponding normalization factor.

#### Receiver operating characteristic (ROC) analysis

For tissue samples, receiver operating characteristic (ROC) curves were used to set the TMT ratio cutoffs that would maximize the retention of true positives, while minimizing the retention of false positives in each case, as described by the Ting lab.^4,5^

Proteins were labeled as true positive based on subcellular location annotations retrieved from uniprot.org. Proteins with any of the following annotations were labeled as true positive (case-insensitive): “secreted”, “endoplasmic reticulum”, “rough endoplasmic reticulum”. Proteins were labeled as false positive if they were found in the human mitomatrix list from supplementary table S1 from Rhee et al., 2013.^6^ Additionally, proteins with signal peptide annotation were removed from the false positive list. Proteins labeled as both true positive and false positive through this approach were reassigned as true positive.

#### Gene Ontology term enrichment analysis

Gene ontology enrichment analysis was performed on the significant upregulated or downregulated proteins (fold-change > 1.5 and p-value < 0.05) using Fisher’s exact test implemented in GOATOOLS against the Gene Ontology (goslim_generic.obo, downloaded on April 24, 2024). An experiment-specific background (i.e. all proteins quantified in the experiment) was used for all analyses except for the cross-experiment analysis (Figure S2F) where the full mouse proteome was used. GO Biological Process terms were mapped to GOslim terms using the GOATOOLS mapslim.py script. After analysis, terms with a Benjamini-Hochberg corrected p-value less than 0.05 were retained.

#### Absolute plasma concentrations

Concentration of plasma proteins quantified by multiple reaction monitoring mass spectrometry in Michaud et *al*. (2018)^7^ were compared with proteins detected in the present study. Of the 385 target analytes detected in C56BL/6J or C57BL/6BR mice in supplementary tables 4, 6, and 7 of Michaud *et al*. (2018), 243 were detected in one or more of the basal, negative energy balance, and obesity conditions. Values were displayed for comparison in Figures 3, S3, 6, and S6, and are available in Tables S2 and S3.

#### UK Biobank

UK Biobank protein-disease associations for prevalent diseases were retrieved from the protein phenome atlas^8^ (proteome-phenome-atlas.com). P-values for diseases were corrected through the Bonferroni correction and only significant associations (Bonferroni adjusted p < 0.05) are shown. Proteins with significant disease associations in UKBB chapters I, IV or IX orthologous to TurboID plasma proteins significantly regulated by low- or high-energy balance are listed in Table S4.

### (F) KEY RESOURCES TABLE

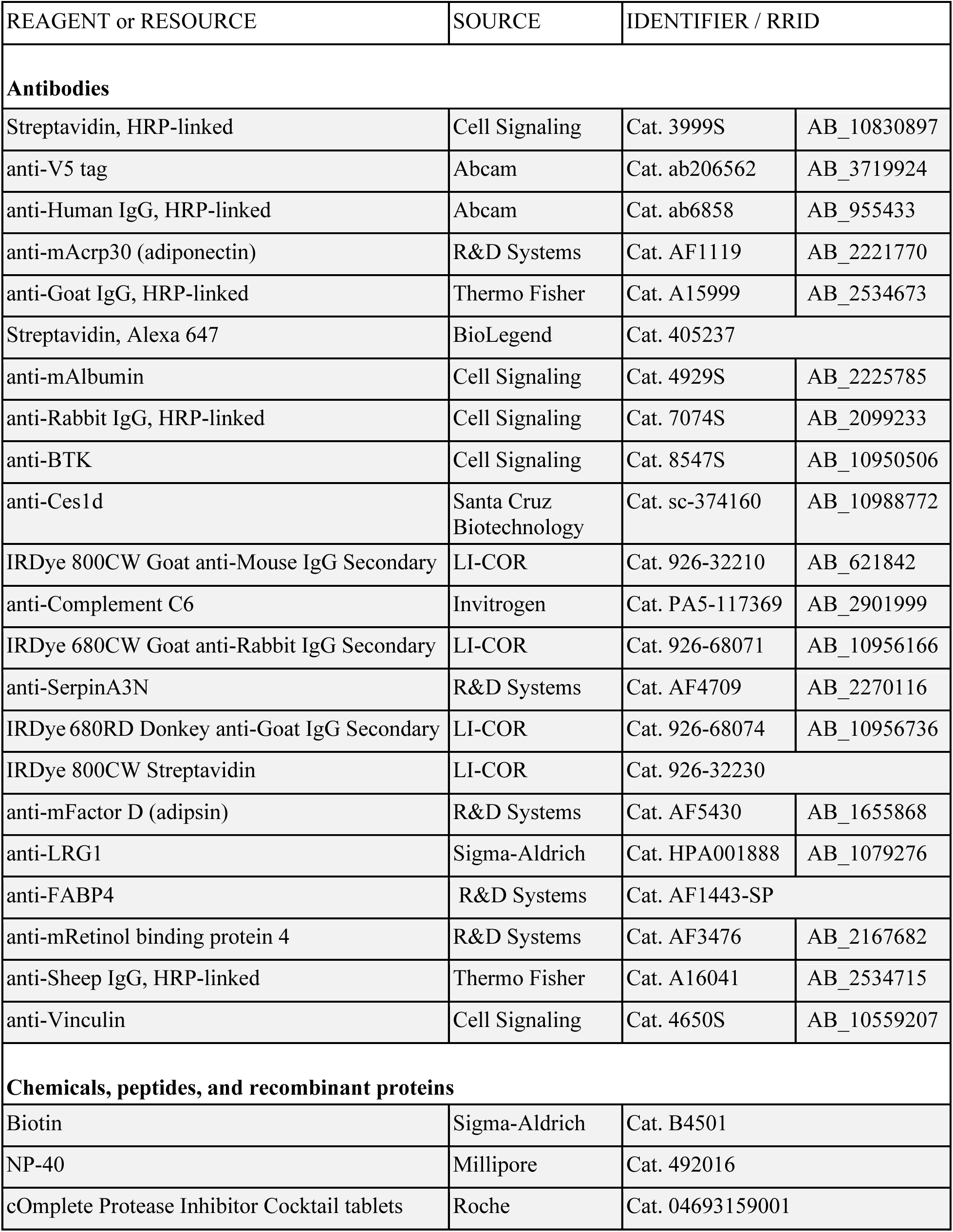
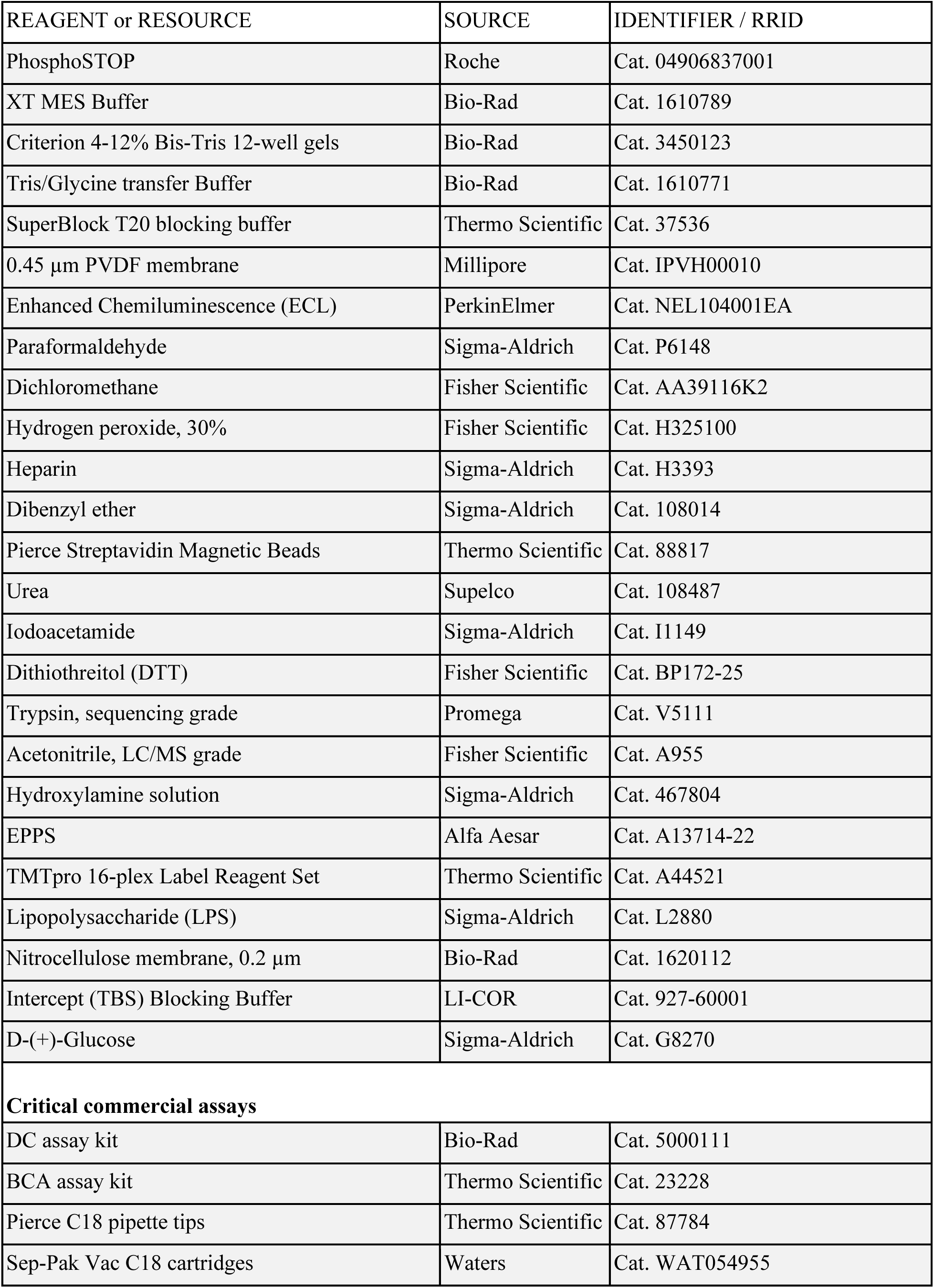
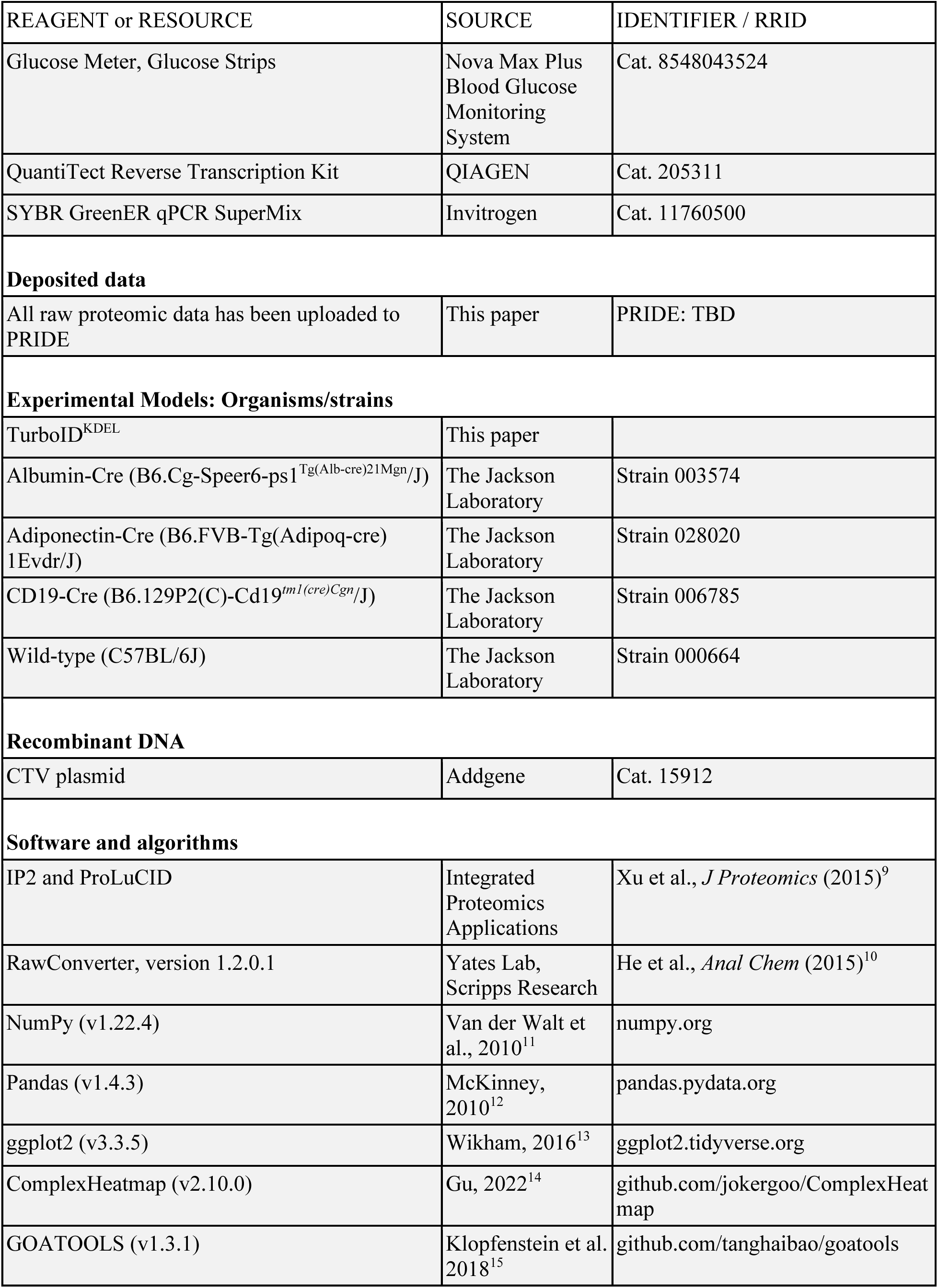
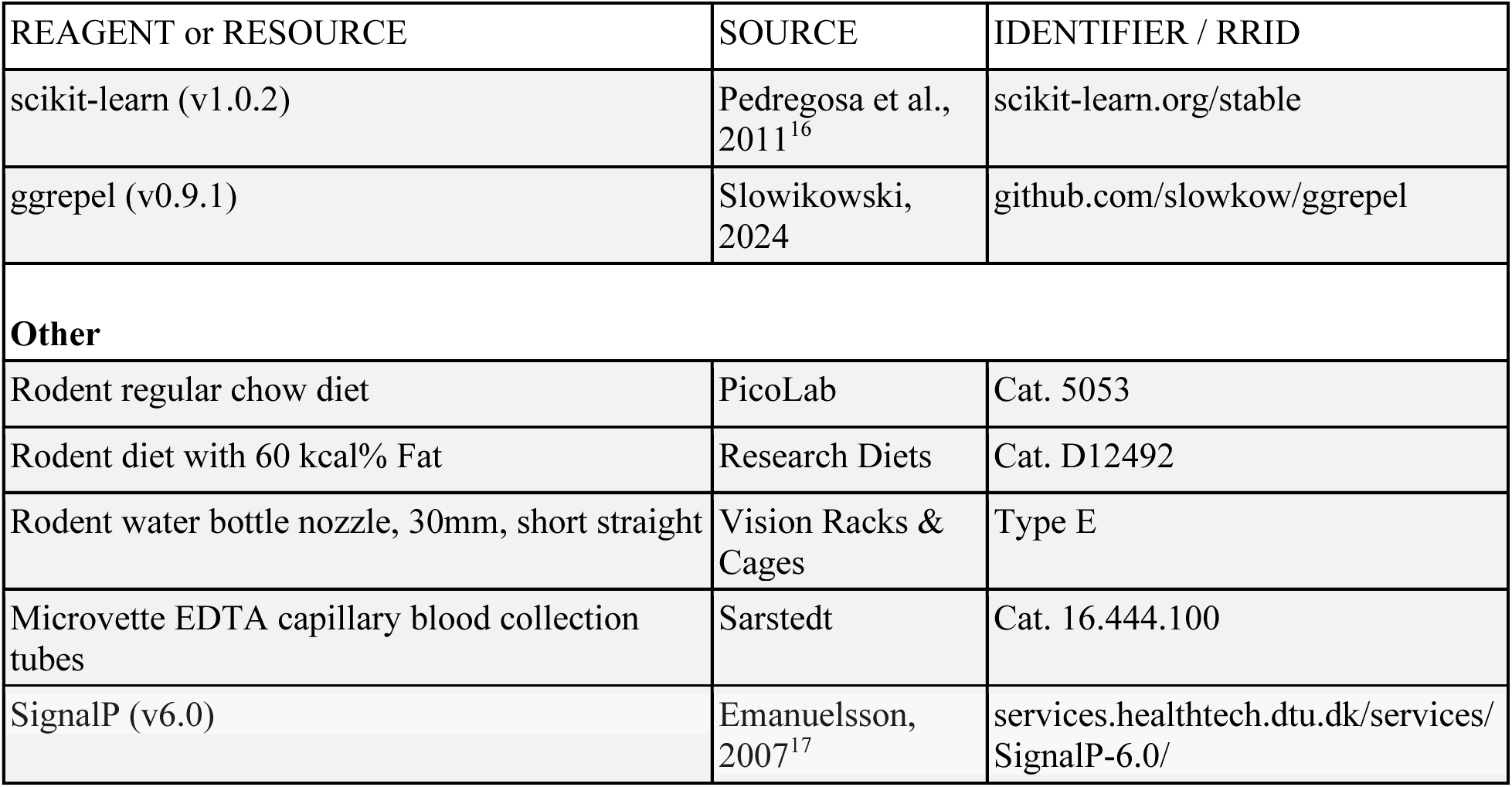

**[F] KEY RESOURCES TABLE**
| REAGENT or RESOURCE | SOURCE | IDENTIFIER / RRID |  |
| --- | --- | --- | --- |
| **Antibodies** |  |  |  |
| Streptavidin, HRP-linked | Cell Signaling | Cat. 3999S | AB_10830897 |
| anti-V5 tag | Abcam | Cat. ab206562 | AB_3719924 |
| anti-Human IgG, HRP-linked | Abcam | Cat. ab6858 | AB_955433 |
| anti-mAcrp30 (adiponectin) | R&D Systems | Cat. AF1119 | AB_2221770 |
| anti-Goat IgG, HRP-linked | Thermo Fisher | Cat. A15999 | AB_2534673 |
| Streptavidin, Alexa 647 | BioLegend | Cat. 405237 |  |
| anti-mAlbumin | Cell Signaling | Cat. 4929S | AB_2225785 |
| anti-Rabbit IgG, HRP-linked | Cell Signaling | Cat. 7074S | AB_2099233 |
| anti-BTK | Cell Signaling | Cat. 8547S | AB_10950506 |
| anti-Ces1d | Santa Cruz Biotechnology | Cat. sc-374160 | AB_10988772 |
| IRDye 800CW Goat anti-Mouse IgG Secondary | LI-COR | Cat. 926-32210 | AB_621842 |
| anti-Complement C6 | Invitrogen | Cat. PA5-117369 | AB_2901999 |
| IRDye 680CW Goat anti-Rabbit IgG Secondary | LI-COR | Cat. 926-68071 | AB_10956166 |
| anti-SerpinA3N | R&D Systems | Cat. AF4709 | AB_2270116 |
| IRDye 680RD Donkey anti-Goat IgG Secondary | LI-COR | Cat. 926-68074 | AB_10956736 |
| IRDye 800CW Streptavidin | LI-COR | Cat. 926-32230 |  |
| anti-mFactor D (adipsin) | R&D Systems | Cat. AF5430 | AB_1655868 |
| anti-LRG1 | Sigma-Aldrich | Cat. HPA001888 | AB_1079276 |
| anti-FABP4 | R&D Systems | Cat. AF1443-SP |  |
| anti-mRetinol binding protein 4 | R&D Systems | Cat. AF3476 | AB_2167682 |
| anti-Sheep IgG, HRP-linked | Thermo Fisher | Cat. A16041 | AB_2534715 |
| anti-Vinculin | Cell Signaling | Cat. 4650S | AB_10559207 |
| **Chemicals, peptides, and recombinant proteins** |  |  |  |
| Biotin | Sigma-Aldrich | Cat. B4501 |  |
| NP-40 | Millipore | Cat. 492016 |  |
| cOmplete Protease Inhibitor Cocktail tablets | Roche | Cat. 04693159001 |  |

| REAGENT or RESOURCE | SOURCE | IDENTIFIER / RRID |
| --- | --- | --- |
| PhosphoSTOP | Roche | Cat. 04906837001 |
| XT MES Buffer | Bio-Rad | Cat. 1610789 |
| Criterion 4-12% Bis-Tris 12-well gels | Bio-Rad | Cat. 3450123 |
| Tris/Glycine transfer Buffer | Bio-Rad | Cat. 1610771 |
| SuperBlock T20 blocking buffer | Thermo Scientific | Cat. 37536 |
| 0.45 µm PVDF membrane | Millipore | Cat. IPVH00010 |
| Enhanced Chemiluminescence (ECL) | PerkinElmer | Cat. NEL104001EA |
| Paraformaldehyde | Sigma-Aldrich | Cat. P6148 |
| Dichloromethane | Fisher Scientific | Cat. AA39116K2 |
| Hydrogen peroxide, 30% | Fisher Scientific | Cat. H325100 |
| Heparin | Sigma-Aldrich | Cat. H3393 |
| Dibenzyl ether | Sigma-Aldrich | Cat. 108014 |
| Pierce Streptavidin Magnetic Beads | Thermo Scientific | Cat. 88817 |
| Urea | Supelco | Cat. 108487 |
| Iodoacetamide | Sigma-Aldrich | Cat. I1149 |
| Dithiothreitol (DTT) | Fisher Scientific | Cat. BP172-25 |
| Trypsin, sequencing grade | Promega | Cat. V5111 |
| Acetonitrile, LC/MS grade | Fisher Scientific | Cat. A955 |
| Hydroxylamine solution | Sigma-Aldrich | Cat. 467804 |
| EPPS | Alfa Aesar | Cat. A13714-22 |
| TMTpro 16-plex Label Reagent Set | Thermo Scientific | Cat. A44521 |
| Lipopolysaccharide (LPS) | Sigma-Aldrich | Cat. L2880 |
| Nitrocellulose membrane, 0.2 µm | Bio-Rad | Cat. 1620112 |
| Intercept (TBS) Blocking Buffer | LI-COR | Cat. 927-60001 |
| D-(+)-Glucose | Sigma-Aldrich | Cat. G8270 |
| **Critical commercial assays** |  |  |
| DC assay kit | Bio-Rad | Cat. 5000111 |
| BCA assay kit | Thermo Scientific | Cat. 23228 |
| Pierce C18 pipette tips | Thermo Scientific | Cat. 87784 |
| Sep-Pak Vac C18 cartridges | Waters | Cat. WAT054955 |
| Glucose Meter, Glucose Strips | Nova Max Plus Blood Glucose Monitoring System | Cat. 8548043524 |
| QuantiTect Reverse Transcription Kit | QIAGEN | Cat. 205311 |
| SYBR GreenER qPCR SuperMix | Invitrogen | Cat. 11760500 |
| **Deposited data** |  |  |
| All raw proteomic data has been uploaded to PRIDE | This paper | PRIDE: TBD |
| **Experimental Models: Organisms/strains** |  |  |
| TurboID<sup>KDEL</sup> | This paper |  |
| Albumin-Cre (B6.Cg-Speer6-ps1<sup>Tg(Alb-cre)21Mgn/J</sup>) | The Jackson Laboratory | Strain 003574 |
| Adiponectin-Cre (B6.FVB-Tg(Adipoq-cre)1Evdr/J) | The Jackson Laboratory | Strain 028020 |
| CD19-Cre (B6.129P2(C)-Cd19<sup>tm1(cre)Cgn/J</sup>) | The Jackson Laboratory | Strain 006785 |
| Wild-type (C57BL/6J) | The Jackson Laboratory | Strain 000664 |
| **Recombinant DNA** |  |  |
| CTV plasmid | Addgene | Cat. 15912 |
| **Software and algorithms** |  |  |
| IP2 and ProLuCID | Integrated Proteomics Applications | Xu et al., *J Proteomics* (2015)<sup>9</sup> |
| RawConverter, version 1.2.0.1 | Yates Lab, Scripps Research | He et al., *Anal Chem* (2015)<sup>10</sup> |
| NumPy (v1.22.4) | Van der Walt et al., 2010<sup>11</sup> | numpy.org |
| Pandas (v1.4.3) | McKinney, 2010<sup>12</sup> | pandas.pydata.org |
| ggplot2 (v3.3.5) | Wikham, 2016<sup>13</sup> | ggplot2.tidyverse.org |
| ComplexHeatmap (v2.10.0) | Gu, 2022<sup>14</sup> | github.com/jokergoo/ComplexHeatmap |
| GOATOOLS (v1.3.1) | Klopfenstein et al. 2018<sup>15</sup> | github.com/tanghaibao/goatools |
| scikit-learn (v1.0.2) | Pedregosa et al., 2011<sup>16</sup> | scikit-learn.org/stable |
| ggrepel (v0.9.1) | Slowikowski, 2024 | github.com/slowkow/ggrepel |
| **Other** |  |  |
| Rodent regular chow diet | PicoLab | Cat. 5053 |
| Rodent diet with 60 kcal% Fat | Research Diets | Cat. D12492 |
| Rodent water bottle nozzle, 30mm, short straight | Vision Racks & Cages | Type E |
| Microvette EDTA capillary blood collection tubes | Sarstedt | Cat. 16.444.100 |
| SignalP (v6.0) | Emanuelsson, 2007<sup>17</sup> | services.healthtech.dtu.dk/services/SignalP-6.0/ |

